# The *HOTAIRM1-miR-222* Axis Regulates Venetoclax Resistance and Defines a High-Risk Subset in Pediatric t(8;21) Acute Myeloid Leukemia

**DOI:** 10.1101/2025.07.12.663834

**Authors:** Christine Wilson, Priyanka Swaroop, Vikas Gaur, Diwakar Sharma, Tanuja Bhardwaj, Sachin Kumar, Anita Chopra, Sampa Ghose, Jayanth Kumar Palanichamy, Deepam Pushpam, Ranjit K. Sahoo, Sameer Bakhshi, Surender K. Sharawat

## Abstract

Although acute myeloid leukemia (AML) with the *RUNX1::RUNX1T1* fusion [t(8;21)(q22;q22.1)] defines a distinct cytogenetic subtype, differences in treatment response suggest additional molecular contributors beyond chromosomal abnormalities. Deregulated hematopoietic lineage-specific long non-coding RNAs (lncRNAs) contribute to leukemogenesis and therapy resistance. To investigate their role in t(8;21) AML, we performed whole-transcriptome sequencing of pediatric patients and age-matched healthy controls, identifying significant downregulation of lncRNA *HOTAIRM1*, a regulator of myeloid differentiation (adjusted P < 0.05). This was confirmed in a single-cell RNA-sequencing dataset (GSE116256) and the Leukemia MILE dataset (GSE13159, P=0.03). Validation of expression in our study cohort using qPCR specifically demonstrated significant downregulation of the myeloid specific isoform, *HOTAIRM1 – HM1V2* (P<0.0001). Analysis of downstream pathways activated by *HM1V2* loss identified *miR-222*, an oncomiR, as a de-repressed target (P=0.01). Elevated *miR-222* expression was observed across AML cell lines (P<0.05), leukemic stem and progenitor cells (GSE117090, P<0.05), AML plasma-derived exosomes (GSE142699, P<0.0001), the current study dataset (P<0.0001), and the TARGET AML dataset (P<0.0001). Restoring *HM1V2* expression with epigenetic agents azacytidine and panobinostat induced apoptosis in venetoclax-resistant Kasumi-1 cells (P < 0.01), through suppression of *miR-222* (P < 0.01) and downregulation of anti-apoptotic proteins BCL-xL and MCL-1 (P < 0.05), key mediators of the venetoclax resistance mechanism. Machine learning based feature selection and Cox regression analysis showed that high *miR-222* expression predicts poor outcome in pediatric t(8;21) AML, validated in both our institutional pediatric AML cohort (P < 0.05) and the multi-institutional TARGET cohort (P < 0.0001). Together, our findings highlight an epigenetic based approach to restore isoform-specific *HM1V2* pathway function in venetoclax-resistant AML cells, and identifies *miR-222* as a prognostic marker to refine risk stratification within the traditionally favorable-risk t(8;21) AML subgroup.

**Key Points:**

1. Loss of myeloid lineage specific isoform of lncRNA *HOTAIRM1* - *HOTAIRM1 variant 2*, results in de-repression of microRNA *miR-222*, and contributes to venetoclax resistance in pediatric AML patients harbouring the t(8;21)(q22;q22.1)/RUNX1::RUNX1T1 fusion.
2. MicroRNA *miR-222* shows potential as a single marker predictor that complements current risk stratification by identifying a subset of pediatric t(8;21) AML patients with poor prognosis.

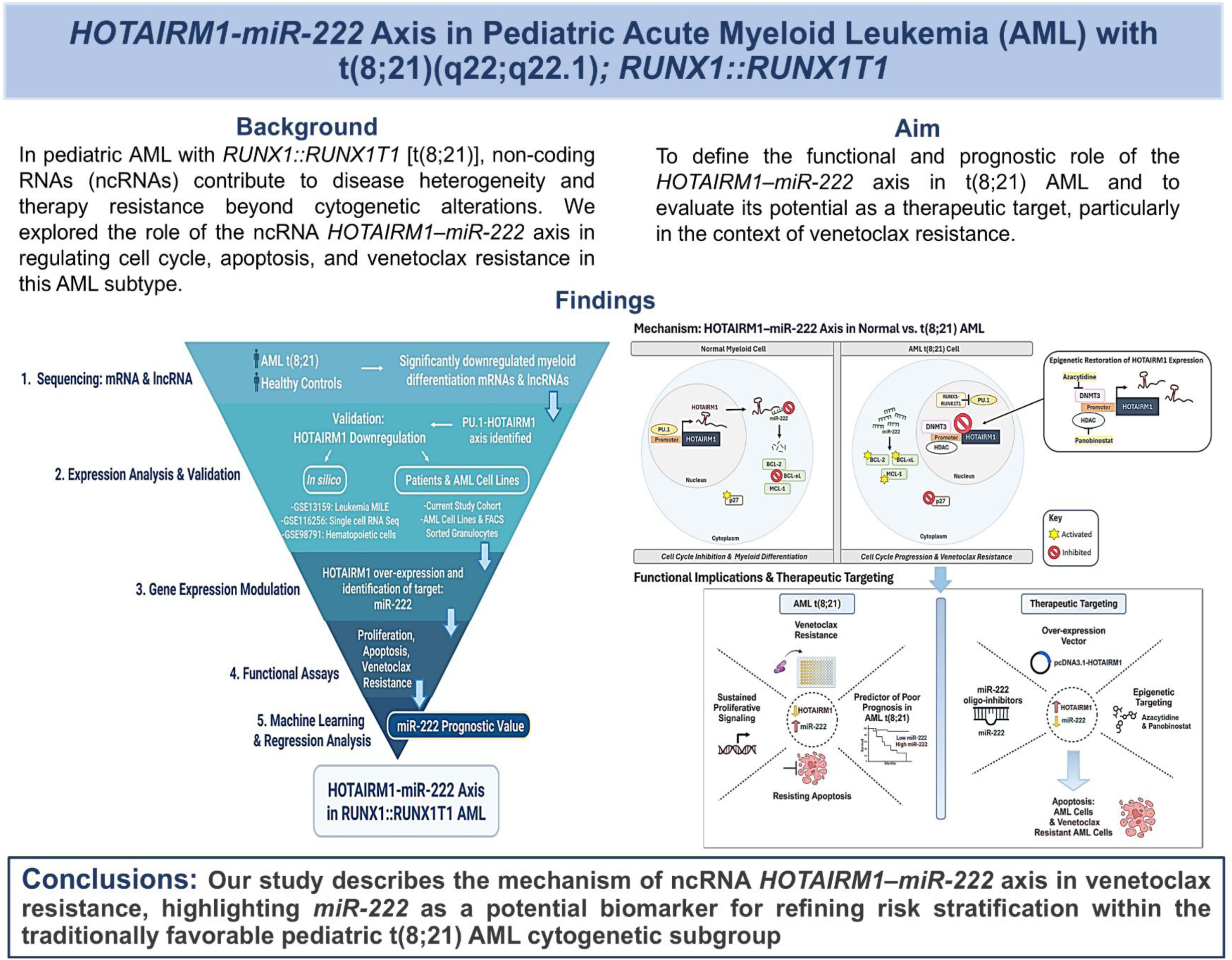

## Introduction

AML with translocation t(8;21)(q22;q22.1) is a cytogenetically defined subgroup traditionally associated with a favorable prognosis, with five-year overall survival rates of around 85% in pediatric patients receiving intensive therapy ^1^. However, despite the high initial response rates, a substantial proportion of pediatric patients i.e. approximately 20–40%, still experience treatment failure or relapse, emphasising the need for a deeper understanding of resistance mechanisms ^2,34,5^. Venetoclax, a selective BCL-2 inhibitor, has been primarily used in relapsed or refractory AML (R/R AML), and is now used in combination with hypomethylating agents (HMAs) for elderly or unfit AML patients ineligible for intensive chemotherapy^6^. More recently, venetoclax has been explored in newly diagnosed AML, particularly in older or chemotherapy ineligible patients, demonstrating improved remission rates and survival ^7^. In pediatric AML, venetoclax-based regimens are showing promising early results^8^. A meta-analysis conducted showed that as a single agent, venetoclax shows limited efficacy in R/R AML, with 72–94% of patients failing to achieve a complete response (CR). However, when combined with HMAs or low-dose cytarabine (LDAC), the refractory disease rates improved, ranging from 43–53% ^9^. Despite the classification of t(8;21) AML as favorable-risk, it is not universally chemosensitive, and responses to venetoclax based regimens in this subgroup have been suboptimal. Yu et al. reported that none of the five relapsed/refractory t(8;21) AML patients achieved remission following venetoclax plus azacitidine therapy ^10^. Similarly, in a separate case series of thirteen treatment-naive t(8;21) AML patients treated with venetoclax in combination with hypomethylating agents, only four achieved complete remission or complete remission with incomplete count recovery (CR/CRi)^11^. Emerging data suggest that additional mutations, such as in the *KIT* gene can drive primary or acquired resistance in t(8;21) AML^12^. These observations emphasise the need to understand the molecular basis of venetoclax resistance in this cytogenetically defined subset, as such insights could refine risk stratification and guide more targeted treatment approaches^13,14^. The lack of response to venetoclax is linked to intrinsic or acquired resistance mechanisms that impair apoptosis. Venetoclax binds to the anti-apoptotic protein BCL-2, displacing pro-apoptotic proteins such as BIM and BAD, and freeing effectors like BAX and BAK to trigger apoptosis. However, resistance to venetoclax develops via upregulation of alternative anti-apoptotic proteins, including BCL-xL and MCL-1, which sustain leukemic cell survival ^15,16^.

Noncoding RNAs, particularly lncRNAs and microRNAs (miRNAs), are important regulators of leukemogenesis and therapeutic resistance. In AML, distinct lncRNA expression patterns are associated with genetic and cytogenetic subtypes, including t(8;21) AML and are linked to differences in disease pathobiology, treatment response, and survival, though their contributions to venetoclax resistance in this context remain underexplored ^17–21^. *HOTAIRM1* is a myeloid differentiation associated lncRNA encoded within the *HOXA* gene cluster and induced by the transcription factor PU.1 during normal haematopoiesis ^22–25^. In AML, *HOTAIRM1* expression is deregulated and the downstream molecular cascade triggered by its aberrant expression is poorly understood ^24, 26–28,29^.

MicroRNAs regulate important cellular processes including proliferation, apoptosis, and drug resistance, by complementary binding to the 3′ untranslated regions (3′ UTRs) of target mRNAs to suppress translation or induce degradation. ^30,31^. MiRNAs, such as *miR-221* and *miR-222*, function as oncomiRs and are linked to poor prognosis in multiple solid cancers^32–34, 35,36, 37^. Despite their known oncogenic roles, the contribution of *miR-221* and *miR-222* to venetoclax resistance in AML has remained unknown.

In this study, we examined the *HOTAIRM1*–*miR-222* regulatory axis in t(8;21) AML aiming to understand its role in leukemic cell survival and therapy resistance. We further assessed its prognostic relevance using Kaplan Meier analysis, multivariable Cox regression, and machine learning based feature selection in both our institutional pediatric AML cohort and the multi-institutional TARGET AML cohort.

## Materials and Methods

### Study Design and Patient Recruitment

A total of 106 patients (≤18 years) with newly diagnosed, *de novo* AML, confirmed by clinical, morphological, immunophenotypic, and cytogenetic criteria, were enrolled at the Dr. B.R. Ambedkar Institute Rotary Cancer Hospital, All India Institute of Medical Sciences (AIIMS), New Delhi, India (i.e. the IRCH cohort). Thirty-six patients were included from an existing cohort (recruited between May, 2018 and September, 2020), and an additional 70 patients were prospectively and consecutively enrolled between September, 2020 and September, 2024. Exclusion criteria included secondary AML, acute promyelocytic leukemia (APL), mixed-phenotype acute leukemia, or granulocytic sarcoma without bone marrow involvement. Age-matched individuals without hematologic malignancies were enrolled as controls. Baseline demographic characteristics, clinical parameters, and European LeukemiaNet (ELN) risk stratification were recorded at diagnosis ^1^. Patient baseline demographics and clinical characteristics have been summarised in Table 1. All patients received standard 3+7 induction as per institutional AML protocol; post-remission, patients underwent either High-Dose Cytarabine (HiDAC) or low-dose cytarabine with daunorubicin. Relapsed cases received Ara-C (cytarabine) + Daunorubicin + Etoposide (ADE) reinduction, which included cytarabine and daunorubicin along with etoposide for 5 days ^38–40^. The study was approved by the Institutional Ethics Committee of AIIMS, New Delhi (IECPG-537/23.09.2020), and conducted in accordance with the Declaration of Helsinki. Written informed consent was obtained from legal guardians, with assent obtained from children aged ≥7 years, as applicable.

**Table 1.**
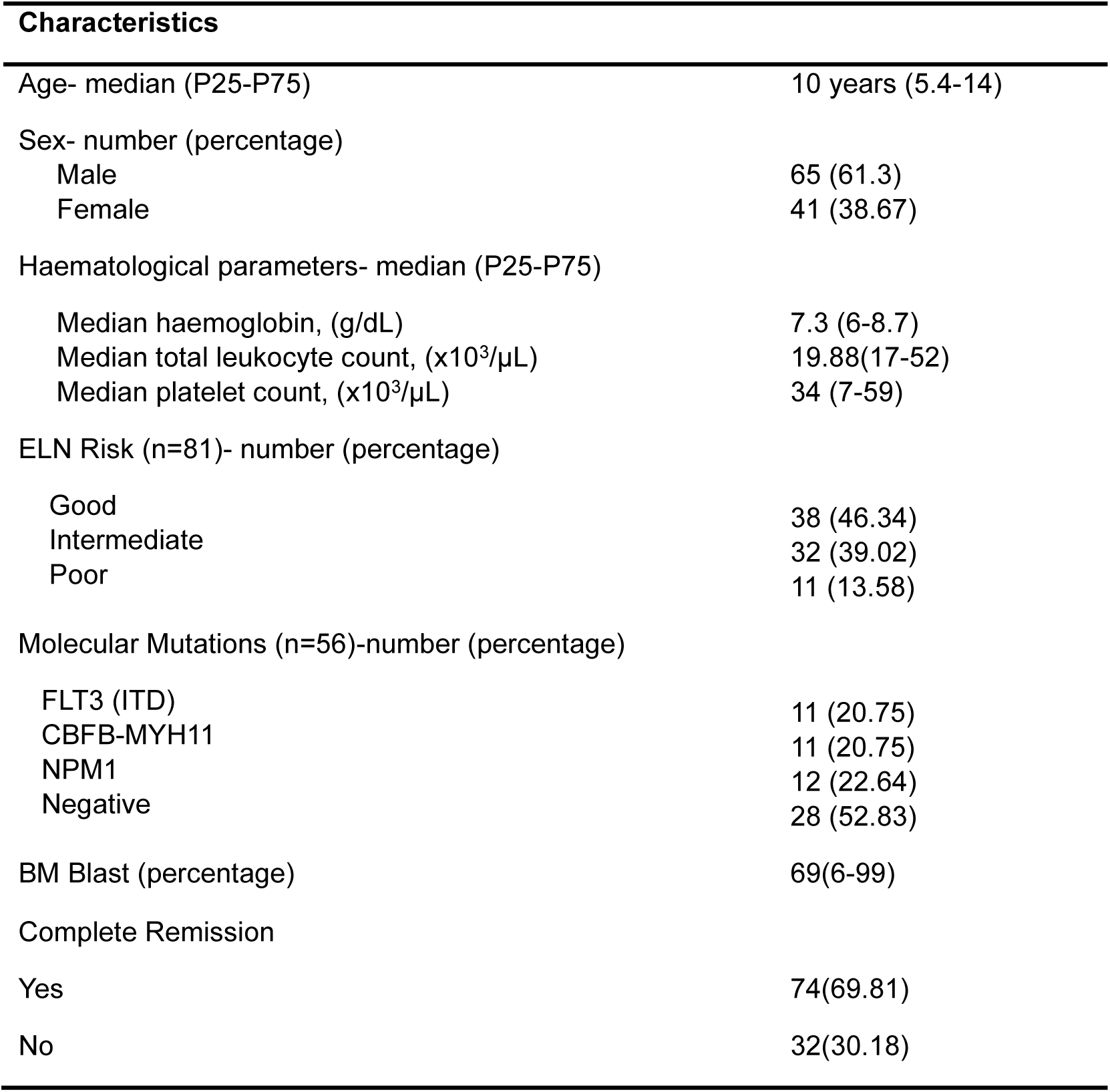
Baseline characteristics of paediatric patients with AML(n=106)

### Whole Transcriptome Sequencing of Patients with t(8;21) AML

RNA isolated from the peripheral blood (PB) samples of six t(8;21) AML patients and five age-matched healthy individuals (RIN >7) were used for library prep (Illumina TruSeq Stranded Total RNA with rRNA depletion) and sequenced on NovaSeq 6000 (150 bp paired-end, ∼100M reads/sample). The corresponding FASTQ files have been deposited in the Sequence Read Archive (SRA) under BioProject ID: PRJNA1279788. Reads were trimmed with fastp and assessed with FastQC/MultiQC (Q20 >95%, Q30 >85%). Trimmed reads were aligned to the GRCh38.p13 reference genome using STAR, with alignment rates exceeding 99% and over 70% of reads mapping uniquely.

Transcripts were assembled with StringTie and annotated with gffcompare. LncRNAs (>200nt, no ORF) were filtered using TransDecoder/CPC2 and classified via UniProt, LNCipedia, GENCODE, and NONCODE. Expression was quantified with featureCounts; transcripts with <5 reads were excluded. Differential expression was assessed using DESeq2, with transcripts considered significant at adjusted p ≤0.05 and absolute log₂ fold change ≥1. Transcripts with fold change ≥2 were prioritized for downstream analysis.

### Retrieval and Analysis of Publicly Available RNA-Sequencing and Micro-array Data of Patients with AML

Gene expression data were retrieved from GEO (GSE98791, GSE116256, GSE117090, GSE209871) using *GEOquery* (v2.60.0) in RStudio (R v4.2.2). Microarray probes were mapped to gene symbols using platform annotations. RNA-seq. data from TARGET-AML and The Cancer Genome Atlas (TCGA)-LAML were obtained via *TCGAbiolinks* (v2.26.0), normalized using *DESeq2* with variance stabilizing transformation. scRNA-seq data (GSE116256) were analyzed using *Seurat* (v4.1.0) with UMAP-based clustering and visualization. A list of all publicly accessed datasets with links have been included in the supplementary table 2.

### *In Silico* MicroRNA Target Prediction

The mature microRNA sequences were retrieved from miRBase (https://www.mirbase.org/) and used to predict targets using four databases-miRDB (http://mirdb.org/), TargetScan (https://www.targetscan.org/), microT-CDS (http://www.microrna.gr/microT-CDS), and TarBase (http://www.microrna.gr/tarbase). Overlapping targets across all databases were shortlisted for further analysis. RNAhybrid (https://bibiserv.cebitec.uni-bielefeld.de/rnahybrid/) was used to assess binding stability to 3′UTRs, with predicted sites showing minimum free energy (mfe) ≤ −20 kcal/mol prioritized.

### Cloning: *HOTAIRM1* for Overexpression Studies and *HOTAIRM1*-miRNA binding Site for Luciferase Reporter Assay

#### Cloning of *HOTAIRM1* into pcDNA3.1+ vector

Full-length *HOTAIRM1* was amplified using Phusion High-Fidelity DNA Polymerase (Thermo Fisher Scientific, Waltham, MA, USA) with primers containing XhoI and EcoRI sites. Amplicons were purified using the QIAquick Gel Extraction Kit (Qiagen, Hilden, Germany), were digested with the respective restriction enzymes (NEB, MA, USA) and ligated with T4 DNA ligase (Thermo Fisher Scientific, Waltham, MA, USA) using standard protocols. The constructs were then transformed into competent DH5α cells. Plasmids were isolated using the QIAprep Spin Miniprep Kit (Qiagen, Hilden, Germany) and verified by restriction digestion and Sanger sequencing (Eurofins Genomics, Bangalore, India). The construct harbouring full length *HOTAIRM1* variant 2 sequence was designated *pcDNA-HM1*, while the empty vector was designated *pcDNA3.1*.

#### Cloning of *HOTAIRM1* and *p27* 3′UTR with *miR-222* MREs into pmirGLO Vector

The *HOTAIRM1* sequence containing *miR-222* microRNA response elements (MREs) and the 3′ untranslated region (3′UTR) of *p27* with *miR-222* MREs were amplified using primers with SalI and SacI sites and cloned into the pmirGLO Dual-Luciferase Reporter Vector (Promega, Madison, WI, USA). Following digestion and ligation using standard protocols as mentioned above, constructs were transformed into competent DH5α, screened, and verified using Sanger sequencing (Eurofins Genomics, Bangalore, India).

### Transient Transfection to Study AML Cell Phenotype after ncRNA Expression Modulation

AML cell lines (THP-1, MOLM-13, Kasumi-1) were transfected with *miR-222* mimics, inhibitors, or scrambled controls (50 nM; Ambion, Thermo Fisher Scientific, Waltham, MA, USA) using Lipofectamine™ 3000 in Opti-MEM™ Reduced Serum Medium (Thermo Fisher Scientific, Waltham, MA, USA) following the manufacturer’s protocol. The cells (1 × 10⁶/well) were seeded in 6-well plates, incubated for 48 h at 37°C, and collected for RNA isolation and qPCR to assess transfection efficiency. For overexpression studies, 1.5 µg of pcDNA3.1+ harbouring the *HOTAIRM1* sequence (pcDNA-HM1) or empty vector was transfected using Lipofectamine™ 3000 and P3000™ Enhancer Reagent (Thermo Fisher Scientific, Waltham, MA, USA). Overexpression was confirmed by qPCR and flow cytometry.

### Dual Luciferase Assay for Interaction Studies Between lncRNA-miRNA and miRNA-mRNA

The interactions between *HOTAIRM1* and *miR-222*, and *miR-222* and the *p27* 3′UTR, were assessed using the Dual-Luciferase® Reporter Assay System (Promega, Madison, WI, USA) per the manufacturer’s protocol. The pmirGLO vector containing either *HOTAIRM1* or *p27* 3′UTR sequences with *miR-222* response elements, along with empty vectors, were co-transfected into AML cell lines with *miR-222* mimic, inhibitor, or scrambled control (Ambion, Thermo Fisher Scientific, Waltham, MA, USA). After 48 h, cells were lysed in Passive Lysis Buffer (Promega), and luminescence was measured using a Dual-Glo® Luminescence Reader (Promega). Firefly luciferase activity was measured first, followed by Renilla luciferase using Stop & Glo® Reagent. Relative luciferase activity was calculated by normalizing Firefly to Renilla signals.

### Fluorescence Activated Cell Sorting (FACS) and Analysis

Cell cycle and apoptosis were assessed using a BD FACS Melody™ flow cytometer (BD Biosciences, San Jose, CA, USA). AML cell lines (Kasumi-1, MOLM-13, THP-1) were treated, fixed in 70% ethanol at 4°C for 30 min, and stained with 50 µg/mL 7-AAD (BD Biosciences) following incubation in binding buffer (0.1% Triton X-100, 0.1% citrate in PBS) at 37°C for 30 min. For apoptosis analysis, cells were stained with 2.5 µL each of Annexin V-PE and 7-AAD using the 7-AAD/Annexin V-FITC Apoptosis Detection Kit (Elabscience, Houston, TX, USA) and incubated in the dark for 15 min. Samples were acquired on the BD FACS Melody™ and analyzed using FlowJo v10.8 (BD Biosciences). Cell cycle phases (G0/G1, S, G2/M) and apoptotic populations (viable, early/late apoptotic, necrotic) were determined based on DNA content and staining patterns. Granulocytes were sorted from normal bone marrow using the BD FACS Melody™ cell sorter (BD Biosciences, San Jose, CA, USA). Bone marrow aspirates were processed to single-cell suspensions following red blood cell lysis (RBC Lysis Buffer). Cells were stained with CD45-FITC (BD Biosciences) and gated based on FSC, SSC, and CD45 positivity. Post-sort analysis confirmed >90% purity. Sorted cells were resuspended in TRIzol® Reagent (Thermo Fisher Scientific, Waltham, MA, USA) for RNA extraction.

### Development and characterisation of venetoclax resistant Kasumi-1 cell line

Venetoclax-resistant Kasumi-1 (Ven-R) cells were generated by gradually exposing parental Kasumi-1 cells to increasing concentrations of venetoclax (MedChemExpress, Monmouth Junction, NJ, USA), starting at the IC20 dose and escalating to IC50, IC70, and IC90. Cells were maintained at each concentration until viability exceeded 90%, assessed via MTT assay. IC50 values were calculated using non-linear regression in GraphPad Prism v9.0 (GraphPad Software, San Diego, CA, USA). Resistance was confirmed by a 7.12-fold increase in IC50 relative to parental cells. Morphological changes were monitored using a brightfield microscope (Magnus Analytics Pvt. Ltd., India). Ven-R cells were further characterized by qPCR analysis of anti-apoptotic genes (*BCL-2, BCL-xL, MCL-1*) and protein expression profiling via flow cytometry using gene-specific antibodies.

### Statistical Analysis

Statistical analyses were performed using RStudio (R v4.2.0) and GraphPad Prism (v9.0). Group comparisons were conducted using chi-square tests for categorical variables and t-tests or Mann–Whitney U tests for continuous variables, as appropriate. One-way ANOVA with Tukey’s post hoc test, or Kruskal–Wallis test with Dunn’s post hoc test, as appropriate was used for multi-group comparisons. Data are reported as mean ± standard deviation (SD) or median with interquartile range (IQR). Event-free survival (EFS) and overall survival (OS) were estimated using the Kaplan– Meier method, with comparisons between groups made using the log-rank test. EFS was defined as the time from registration to the first event (induction failure, relapse, or death), with patients without an event censored at last follow-up. OS was defined as the time from the date of registration to death from any cause, with patients alive at last follow-up as censored date. Cox proportional hazards models were used for univariable and multivariable analysis, adjusting for age, gender, cytogenetic risk, and blast percentage. Forest plots were generated using the survival and forestplot R packages in R Studio. Machine learning based feature selection was conducted using the randomForest R package. Variables included clinical parameters and miRNA expression; importance was ranked by mean decrease in Gini index. For drug combination studies, the Chou–Talalay method was used to calculate the combination index (CI), with CI <1 indicating synergy. P-values <0.05 were considered significant. Experiments were conducted in triplicate unless stated otherwise.

## Results

### Dysregulation of Myeloid Differentiation Genes Highlights Repression of the *PU.1*-*HOTAIRM1* Pathway in t(8;21) AML

To identify important regulators of myeloid differentiation that are silenced in t(8;21) AML, we performed RNA sequencing on a discovery cohort of patients with t(8;21) AML (n=6) and healthy donors (n=5) (figure 1A). Differential expression analysis identified a distinct subset of significantly downregulated mRNAs and lncRNAs in t(8;21) AML (figure 1B). We focused on the downregulated mRNAs associated with the gene ontology (GO) term “myeloid cell differentiation” (GO:0002521). In t(8;21) AML, eleven genes involved in myeloid lineage commitment were silenced (figure 1C). To extend this analysis to the non-coding transcriptome, we referenced a dataset from Schwarzer et al., 2017 to identify lncRNAs enriched in normal myeloid differentiation^20^. Cross-referencing this dataset with our differential expression results, we identified two lncRNAs (*HOTAIRM1* and *LINC00173*), which specifically regulate myeloid differentiation and serve as myeloid fingerprint lncRNAs, both of which were significantly downregulated in t(8;21) AML (figure 1D).

**Figure 1.**
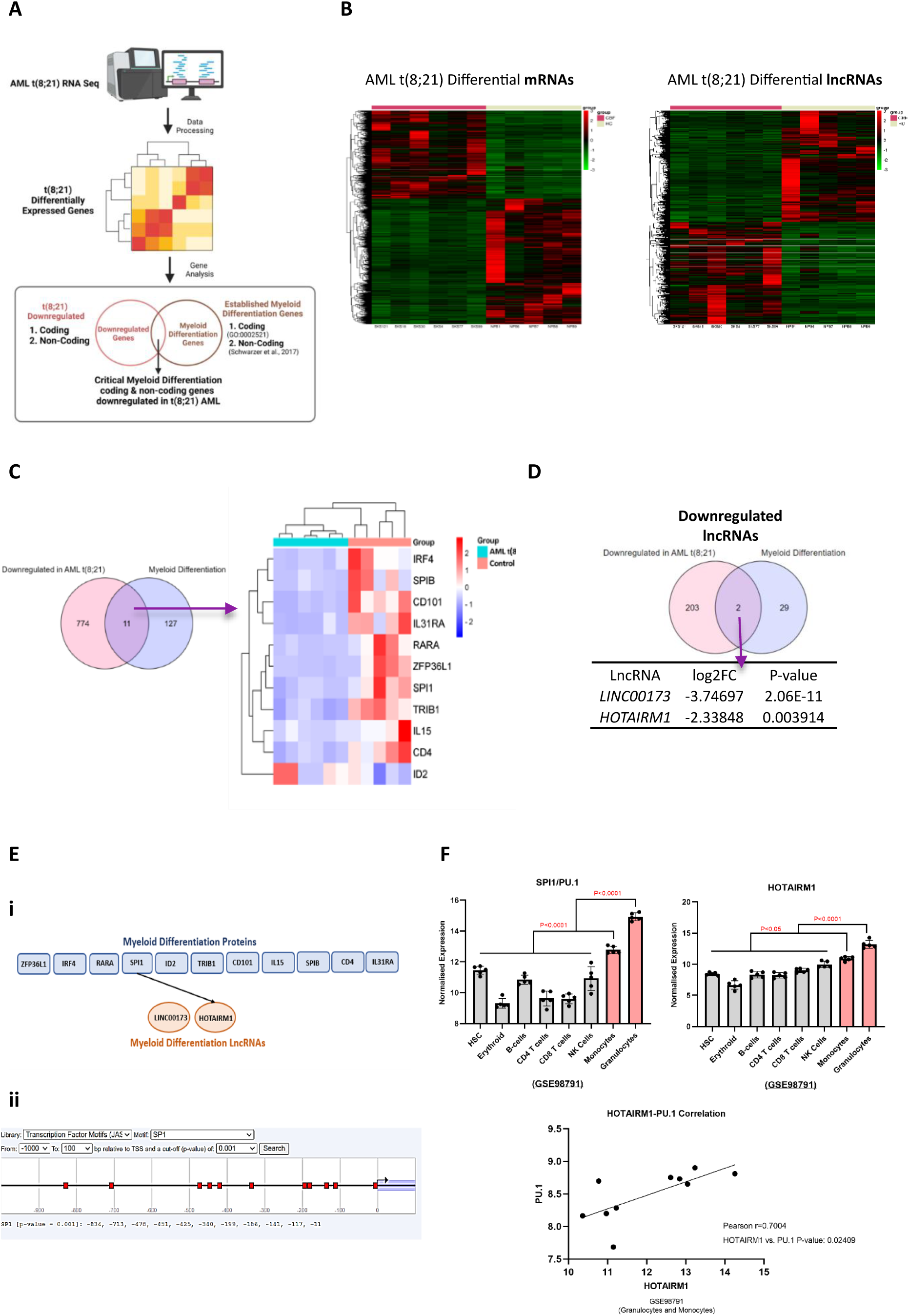
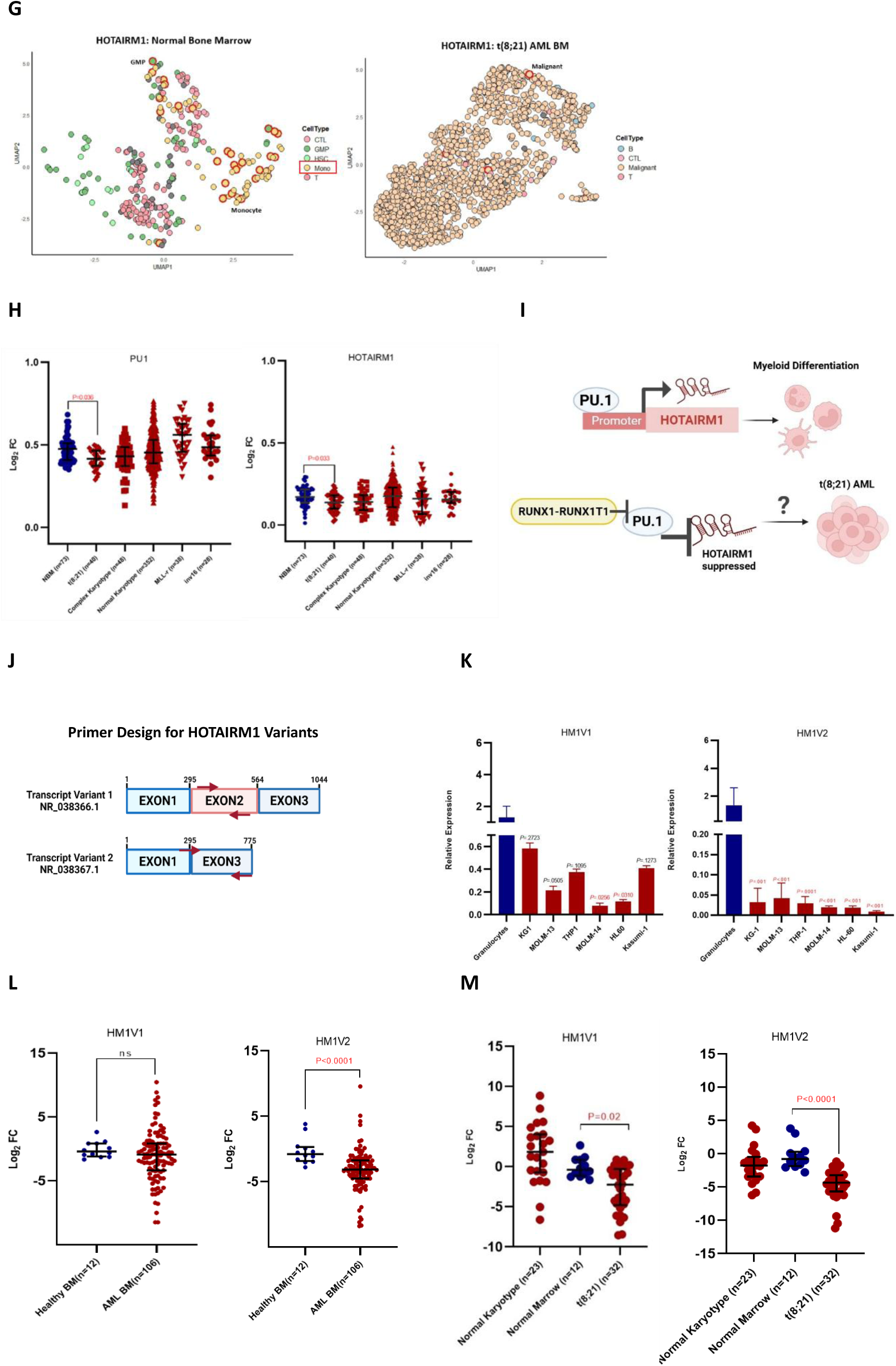
LncRNA *HOTAIRM1*, downstream of myeloid transcription factor PU.1 is enriched in differentiated myeloid cells and downregulated in t(8;21) AML. **A.** Schematic: Identification of myeloid differentiation coding and non-coding genes downregulated in t(8;21) AML. **B.** Heatmap showing differential expression of mRNAs and lncRNAs between AML t(8;21) and healthy controls. **C.** Venn diagram and heatmap depicting 11 coding genes associated with myeloid differentiation (GO:0002521), that are significantly downregulated in AML. **D.** Venn diagram and table listing 2 significantly downregulated lncRNAs in t(8;21) AML which are critical for myeloid differentiation i.e. *LINC00173* and *HOTAIRM1*. **E. i.** Schematic depicting relationship between t(8;21) AML downregulated coding and non-coding myeloid differentiation genes. Evidence in the literature indicates that SPI1 protein (also known as PU.1) directly binds to the promoter of *HOTAIRM1* and induces its expression. **ii.** *HOTAIRM1* promoter was analysed for transcription factor motifs using the JASPAR database and 11 binding sites for SPI1/PU.1 were found (P=0.001). **F.** Expression analysis of *SPI1/PU.1* and *HOTAIRM1* in different hematopoietic cell populations from the <u>GSE98791</u> dataset. *PU.1* and *HOTAIRM1* were both significantly upregulated in monocytes(P<0.0001) and granulocytes(P<0.0001) relative to other hematopoietic cells, and their expression showed a strong positive correlation (r=0.7, P=0.02). **G.** UMAP plot visualizing single-cell RNA-seq data of bone marrow samples from healthy individuals and an AML t(8;21) patient (<u>GSE116256</u>), highlighting cell types expressing *HOTAIRM1* (cells encircled in red). Cells with a thick red border express *HOTAIRM1*. *HOTAIRM1* is enriched in monocytes from healthy bone marrow, however negligible expression is observed in t(8;21) marrow. **H.** PU.1 and *HOTAIRM1* expression levels were plotted from the leukemia MILE study (<u>GSE13159</u>) in AML subtypes and healthy controls. Patients harbouring the t(8;21) translocation have significantly lower levels of PU.1 (P=0.036) and *HOTAIRM1* (P=0.033). **I.** Hypothesis: PU.1 normally binds to HOTAIRM1 and promotes its expression, facilitating normal myeloid differentiation. However, when PU.1 is sequestered by the oncoprotein RUNX1-RUNX1T1 and *HOTAIRM1* is not induced, does this disruption trigger leukemogenic pathways that contribute to the development of t(8;21) AML? **J.** Two transcript variants of *HOTAIRM1* are reported in NCBI. *HOTAIRM1 transcript variant 1* (NR_038366.1) has 3 exons and length of 1044 base pairs, whereas a shorter variant i.e. *HOTAIRM1 transcript variant 2,* also exists (NR_038367.1) which lacks EXON2 and is of size 775 base pairs. **K.** *HOTAIRM1* was quantified using qPCR in 6 AML cels lines along with granulocytes sorted from bone marrow of healthy individuals using FACS. The expression of *HM1V1* is significantly lower in MOLM-13 (P=0.0256) and HL-60 (P=0.0310) AML cell lines, whereas *HM1V2* is downregulated in all AML cell lines, relative to granulocytes (KG-1, MOLM-14, HL-60, Kasumi-1, MOLM-13, THP-1, P<0.05). **L.** Expression analysis of *HOTAIRM1* in IRCH cohort: *HM1V1* is not significantly different in patients with AML relative to normal marrow, whereas *HM1V2* is downregulated in patients with AML relative to normal marrow (P=0.0006). **M.** Analysis of AML sub-groups from the IRCH cohort based on chromosomal translocations indicates that both *HM1V1* and *HM1V2* are significantly downregulated in patients with the t(8;21) translocation (P<0.0001). Statistical analyses across datasets were performed using unpaired t-tests or one-way ANOVA for parametric data, and Mann–Whitney U test or Kruskal–Wallis test for non-parametric data, with appropriate post hoc corrections. P-values <0.05 were considered statistically significant.

The schematic in figure 1E(i) illustrates the regulatory relationship between downregulated myeloid differentiation protein-coding genes and lncRNAs in t(8;21) AML from our dataset. Among the myeloid differentiation-related genes downregulated in AML (as shown in figure 1C), *SPI1* (encoding *PU.1*) is a central transcription factor known to induce *HOTAIRM1* expression, one of the two significantly repressed lncRNAs identified (figure 1D). Promoter analysis using the Eukaryotic Promoter Database (EPD) revealed 11 putative PU.1 binding motifs within the *HOTAIRM1* promoter region (figure 1E(ii)), supporting its regulation by this myeloid-specific transcription factor. We further assessed the expression of *PU.1* and *HOTAIRM1* in independent transcriptomic datasets. Analysis of the GSE98791 microarray dataset showed enrichment of both *PU.1* and *HOTAIRM1* in mature myeloid cells i.e. monocytes and granulocytes (P < 0.0001 and P = 0.0001, respectively) (figure 1F). Additionally, a strong positive correlation was observed between *HOTAIRM1* and *PU.1* expression within these mature myeloid cell types (Figure 1F, P=0.01, r=0.7). Single-cell RNA-sequencing data from GSE116256 further confirmed this, showing high expression of *HOTAIRM1* in monocytes from healthy marrow (encircled in red), compared to negligible expression in cells from a t(8;21)-positive AML patient (figure 1G). Considering the low expression of *PU.1* and *HOTAIRM1* in t(8;21) AML, we further analysed its expression in patient samples from the Leukemia MILE cohort (GSE13159), where *PU.1* and *HOTAIRM1* were both significantly downregulated in t(8;21) AML samples (n=40) compared to normal bone marrow (n=73) (P = 0.036 and P = 0.033, respectively) (figure 1H). These observations reinforce a model in which RUNX1-RUNX1T1 represses PU.1, thereby indirectly silencing *HOTAIRM1* and impairing normal myeloid differentiation (figure 1I).

The lncRNA *HOTAIRM1* has two transcript variants, a longer, *HOTAIRM1* variant 1 [*HM1V1*; NR_038366.1; 1044bp] and a shorter *HOTAIRM1* variant 2 [*HM1V2*; NR_038367.1; 775bp] - (figure 1J). Across multiple AML cell lines, including the t(8;21)-positive Kasumi-1 cell line, *HM1V2* expression (qPCR) was significantly downregulated compared to granulocytes sorted from healthy individuals (P < 0.05) (figure 1K). However, *HM1V1* showed reduced expression only in MOLM-14 (P=0.02) and HL60 (P=0.03) cell lines. In bone marrow samples from our study cohort, *HM1V2* expression was reduced compared to normal bone marrow (P < 0.0001), while *HM1V1* showed no significant difference between patients with AML and healthy controls (figure 1L). Subtype-specific analysis showed that both isoforms were downregulated in patients with t(8;21) AML, with *HM1V2* showing more evident downregulation (P = 0.02 for *HM1V1*, P < 0.0001 for *HM1V2*) (figure 1M). These findings highlight the coordinated dysregulation of the *PU.1*-*HOTAIRM1* axis in t(8;21) AML.

### Effect of *HM1V2* Overexpression on the t(8;21) subtype of AML and its Downstream Target MicroRNAs

To evaluate the functional relevance of *HOTAIRM1* loss in patients with t(8;21) AML, we used white blood cells (WBCs) isolated from the peripheral blood of a healthy individual to amplify the full-length transcript of *HOTAIRM1* (supplementary figure 1A). The product (775bp) was cloned into the pcDNA3.1+ vector and confirmed as *HM1V2* by Sanger sequencing (supplementary Figure 1A–E). Subsequent functional studies were conducted using the cloned *HIMV2* variant (pcDNA-HM1). The predicted secondary structure of the cloned transcript matched the NCBI reference, with comparable minimum free energy and conserved stem-loop conformations (supplementary figure 1E). Transfection of AML cell lines with pcDNA-HM1 resulted in significant overexpression (qPCR) of *HM1V2* compared to empty-vector controls (pcDNA3.1), in Kasumi-1 (P = 0.01), THP-1 (P = 0.007), and MOLM-13 (P = 0.001) cells (figure 2A). Its corresponding target, CD11b expression was significantly upregulated (flow cytometry) (supplementary Figure 1G). We evaluated the impact of *HM1V2* overexpression on cell cycle progression using 7-AAD based flow cytometry analysis. In t(8;21)-positive Kasumi-1 cells, *HM1V2* induced a G0–G1 arrest (P = 0.0001), However no significant changes in the cell cycle were detected in THP-1 or MOLM-13 cells (figure 2B, supplementary figure 1H), indicating that the effects of *HM1V2* are specific to the t(8;21) subtype of AML. Annexin V/7-AAD staining showed a significant increase in early apoptosis in Kasumi-1 cells overexpressing *HM1V2* (P = 0.0005), an effect absent in MLL-rearranged THP-1 and MOLM-13 cell lines (figure 2C, supplementary Figure 1I). These data suggest a selective pro-apoptotic and cell cycle-regulatory function of *HM1V2* in t(8;21) AML.

**Figure 2.**
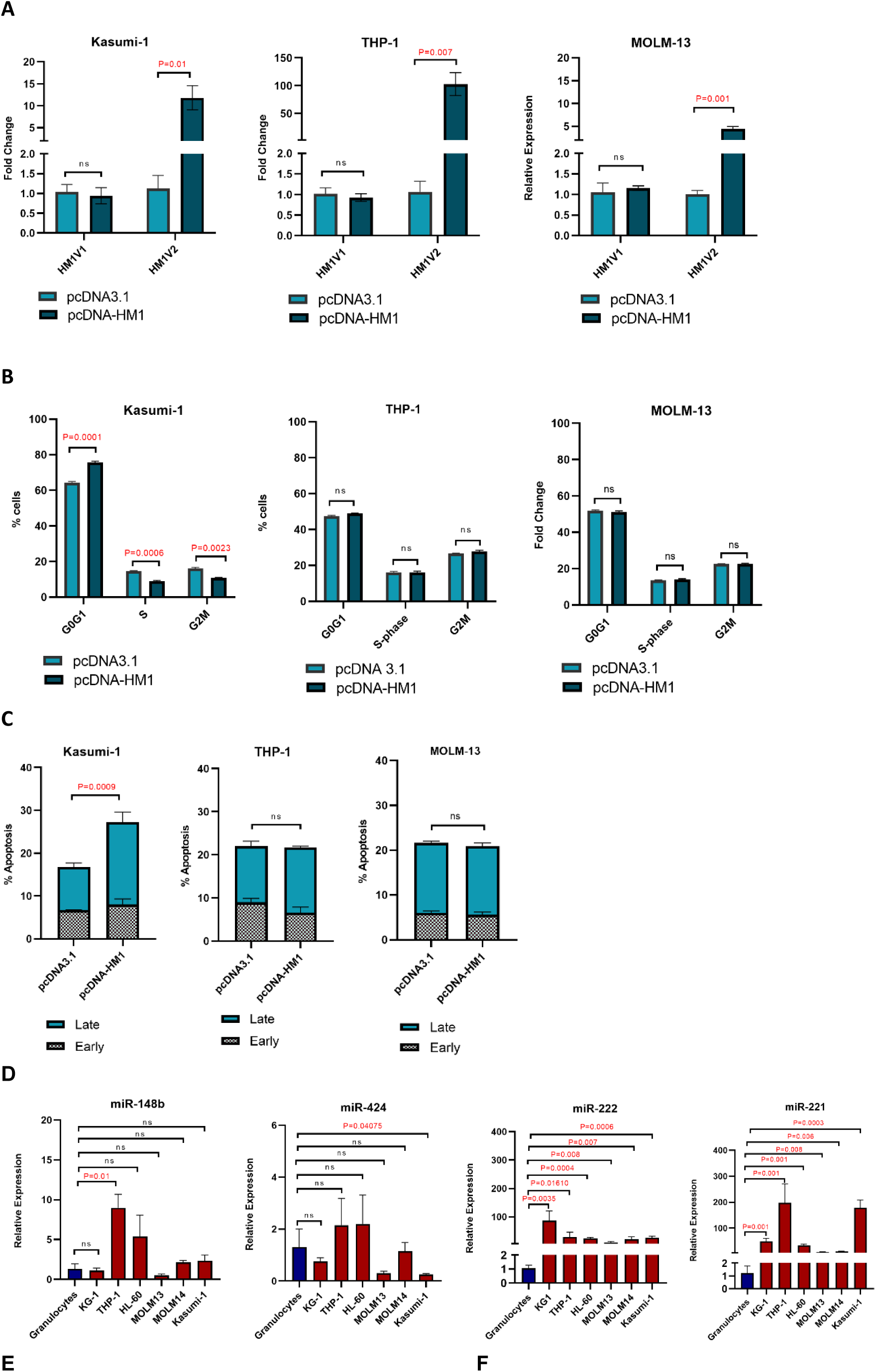
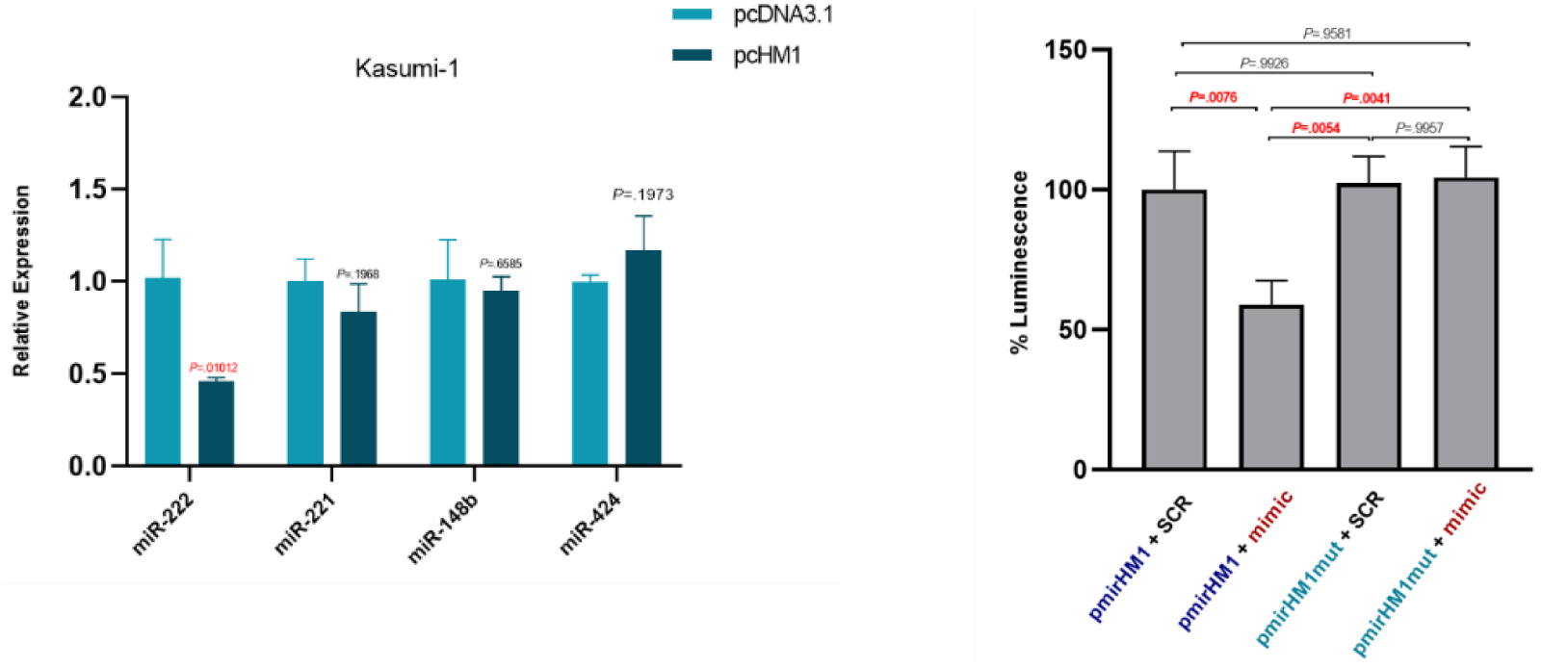
*HOTAIRM1 variant 2* overexpression results in G0G1 arrest and increased apoptosis in t(8;21) AML positive cell line Kasumi-1. A. Expression of *HM1V1* and *HM1V2* in Kasumi-1 (P=0.62, P=0.01796), THP-1(P=0.61, P=0.007), and MOLM-13(P=0.001) after transfection of cells with pcDNA3.1+ vector harbouring full length sequence of *HOTAIRM1*. **B.** Cell cycle analysis of AML cell lines after overexpression shows that *HM1V2* induces G0-G1 arrest in Kasumi-1 (P=0.0001) but not in THP-1 and MOLM-13 cell lines. **C.** The overexpression of *HM1V2* results in significantly high levels of apoptosis in Kasumi-1 (P=0.0005) but has no effect in THP-1 and MOLM-13 cell lines. **D.** Expression of putative microRNA regulators of *HM1V2* in AML cell lines. MicroRNAs *miR-222* and *miR-221* are significantly overexpressed in all AML cells relative to granulocytes isolated from healthy individual (P<0.05). E. *HM1V2* overexpression leads to significant downregulation of microRNA *miR-222* (P=0.01). **F.** Co-transfection of HEK293T cells with the *HM1V2* sequence harbouring the miR-222 binding region, along with a *miR-222* mimic confirmed the direct interaction between the two non-coding RNAs *in vitro* (P=0.007). Statistical analysis was performed using unpaired t-tests or one-way ANOVA for parametric data, followed by Tukey’s post hoc test for multiple group comparisons or Dunnett’s test for comparing multiple groups to a control. P-values <0.05 were considered statistically significant.

We profiled microRNAs (*miR-148b, miR-424, miR-222 and miR-221*) predicted to interact with *HM1V2* (supplementary figure 1J) to study the downstream regulatory effects. Baseline expression analysis revealed consistent and significant upregulation of *miR-222* and *miR-221* in all AML cell lines (figure 2D, P<0.05). The overexpression of *HM1V2* led to target-specific reduction in *miR-222* (figure 2E, P = 0.01). The interaction between *HM1V2* and *miR-222* was further validated by the dual-luciferase reporter assay, confirming a direct in vitro binding between *HM1V2* and *miR-222* (figure 2F, P=0.007).

### MicroRNAs *miR-221* and *miR-222* are upregulated in AML and influence leukemic cell proliferation and apoptosis

*MiR-222* and *miR-221* both share a seed region and belong to the same family of microRNAs. Although *HM1V2* overexpression influenced *miR-222* expression specifically, we studied the expression patterns of both microRNAs in AML, due to their shared regulatory potential (figure 3A). In data from the GSE117090 dataset, *miR-221* was upregulated in leukemic stem cells (LSCs) relative to hematopoietic stem cells (HSCs) (P = 0.0001), and in leukemic progenitor cells (LPCs) relative to hematopoietic progenitor cells (HSPCs) (P = 0.002), while *miR-222* was enriched in LPCs when compared to normal progenitors (P = 0.02) (figure 3B). In addition to qPCR analysis (figure 2D), the GSE51908 dataset confirmed upregulation of both miRNAs in AML cell lines vs. healthy granulocytes (P < 0.05) (supplementary figure 2A). In bone marrow samples from the current study cohort (AML n = 106) both microRNAs were significantly upregulated (*miR-221*, P = 0.001 and *miR-222*, P < 0.0001) compared to controls i.e. normal bone marrow (figure 3C). Cytogenetic subgroup analysis demonstrated significantly higher miR-221 and miR-222 expression in t(8;21) AML (P = 0.007 and P < 0.0001, respectively) and in normal karyotype (NK AML) (P = 0.001 and P < 0.0001, respectively) relative to normal bone marrow (Figure 3D). Further, upregulation of both microRNAs was observed in an independent miR-seq dataset of AML patients that were cytogenetically normal (CN-AML)/ NK AML patients compared to controls i.e. normal bone marrow (P < 0.05) (supplementary figure 2B). In the TARGET cohort (AML, n = 283), both miRNAs were significantly upregulated at diagnosis compared to healthy bone marrow (P < 0.0001), and their expression levels were further elevated in relapsed patients relative to baseline (P < 0.01). Cytogenetic sub-group analysis showed significant upregulation of both *miR-221* (P<0.0001) and *miR-222* (P<0.0001) in t(8;21) AML as well as NK AML. Expression analysis in peripheral blood (P = 0.0005 for *miR-222*; P = 0.0431 for *miR-221*) and plasma-derived exosomes (GSE142699; P < 0.0001) further confirmed their increased expression in AML (supplementary Figure 2C-D). Beyond hematologic malignancies, both *miR-221* and *miR-222* also exhibited high expression across multiple solid tumors, including triple-negative breast cancer (TNBC), estrogen-dependent breast cancer, ovarian cancer, and thyroid cancer (supplementary figure 2E). Analysis of solid tumour datasets from TCGA showed *miR-222* overexpression in 16 of 22 tumor types, (supplementary figure 2F). These above data demonstrate consistent and cytogenetic subtype-independent upregulation of *miR-221* and *miR-222* in AML and across several cancer types.

**Figure 3.**
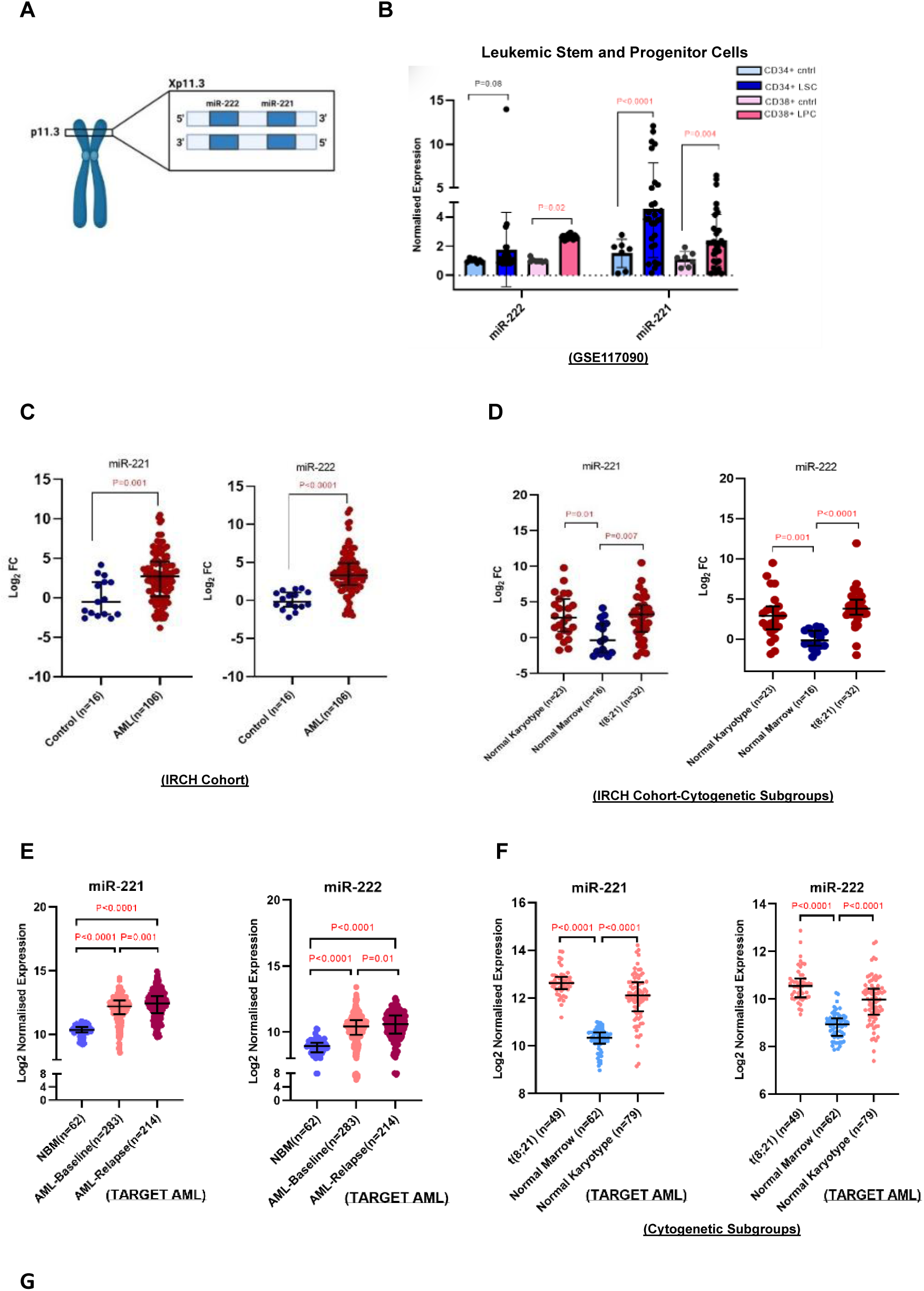
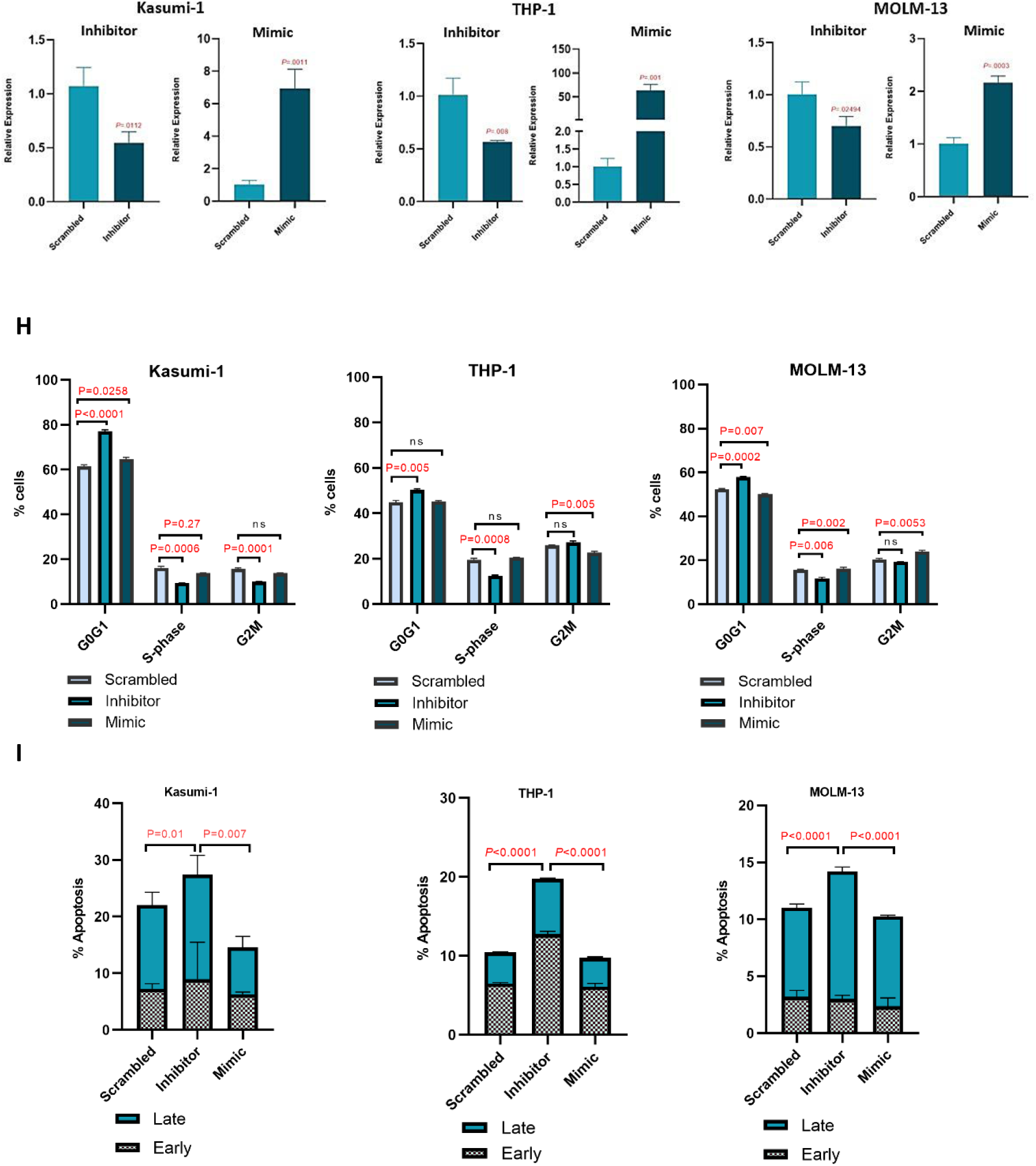
MicroRNAs *miR-221* and *miR-222* are upregulated in AML and influence leukemic cell proliferation and apoptosis. **A.** MicroRNA *miR-221* and *miR-222* are encoded on the X-chromosome and belong to the same family i.e. they harbour the same regulatory seed region. **B.** The <u>GSE117090</u> dataset was utilised to assess expression levels of *miR-221* and *miR-222* in hematopoietic stem and progenitor cells vs. AML LSCs and AML LPCs. AML LSCs and LPCs showed upregulated expression of *miR-221*(P=0.0001), while *miR-222* was enriched in AML LPCs(P=0.02). **C.** Expression of *miR-222* and *miR-221* was quantified in bone marrow samples of patients from the IRCH cohort (n=106) using qPCR. **D.** Subsequently analysis was performed on cytogenetic sub-group, which showed significant upregulation of both *miR-221*(P=0.001) and *miR-222* (P<0.0001), irrespective of cytogenetic sub-group. **E.** Expression of *miR-221* was upregulated in AML samples from the TARGET dataset at baseline compared to healthy marrow (P<0.0001), as well as at relapse compared to baseline (P=0.001). Similarly, *miR-222* was also upregulated in patients at baseline compared to normal (P<0.0001), and at relapse compared to baseline (P=0.001) in the TARGET cohort. **F.** Sub-group analysis of the TARGET cohort showed significant upregulation of *miR-221* (P<0.0001) and *miR-222* (P<0.0001) in t(8;21) AML as well as in patients with normal karyotype (NK-AML). **G**. Transfection efficiency of *miR-222* inhibitor and *miR-222* mimic in Kasumi-1, THP-1 and MOLM-13 leukemic cell lines. The *miR-222* oligo inhibitor leads to significant downregulation of *miR-222* in Kasumi (P=0.0112), THP-1(P=0.008) and MOLM-13 (P=0.02), while *miR-222* mimic leads to signfiicantly high *miR-222* expression in Kasumi-1(P=0.0011), THP-1 (0.001) and MOLM-13 (P=0.0003). **H.** Cell cycle analysis based on 7-AAD staining by flow cytometry showed there was significant G0-G1 phase arrest in Kasumi-1(P<0.0001), THP-1(P=0.001) and MOLM-13(P=0.0001) when treated with the inhibitor for *miR-222*. **I.** The downregulation of *miR-222* led to significant increase in the percentage of apoptosis in Kasumi-1, THP-1 and MOLM-13 (P<0.0001). Statistical analyses were conducted using Student’s t-test or one-way ANOVA for parametric data, with Tukey’s/Dunnett’s post hoc test applied for multiple group comparisons. Non-parametric data were analyzed using the Mann–Whitney U test or Kruskal– Wallis test followed by Dunn’s multiple comparisons test. P-values <0.05 were considered statistically significant.

Considering the regulation of *miR-*222 by *HM1V2,* we next performed gain and loss-of-function assays to study the molecular role of this oncomiR. Transfection with *miR-222* inhibitors led to a significant reduction in its expression (P < 0.05), while mimic transfection resulted in overexpression (P < 0.01) across Kasumi-1, THP-1, and MOLM-13 cell lines (figure 3G). Cell cycle analysis revealed G0-G1 arrest upon *miR-222* inhibition (P < 0.0001), with reduced S-phase progression in all cell lines (P ≤ 0.001) (figure 3H). Further assessment of apoptosis using Annexin V/7-AAD staining demonstrated that *miR-222* inhibition significantly increased the proportion of apoptotic cells across all three AML cell lines i.e. Kasumi-1 (P < 0.0001), THP-1 (P < 0.0001), and MOLM-13 (P < 0.0001) (figure 3I).

### *miR-222* negatively regulates the cell cycle inhibitor CDKN1B (p27^Kip^^1^) and promotes venetoclax resistance in t(8;21) AML

To explore the functional role of *miR-222* in regulating cell cycle and apoptosis pathways, we first identified putative targets using four miRNA prediction databases (miRDB, TargetScan, mR-microT, and TarBase). CDKN1B (p27^Kip1^), a known G1-S checkpoint regulator and tumor suppressor, was a common predicted target among the four databases ^41^ (figure 4A). RNAHybrid analysis identified two *miR-222* MREs in the 3′UTR of p27^Kip1^: MRE (1) (8-mer, −25.3 kcal/mol) and MRE (2) (7-mer, −21.5 kcal/mol) (figure 4B). To validate this interaction, luciferase assays were performed in HEK293T cells co-transfected with *miR-222* mimics and a luciferase reporter construct containing the *p27* 3′UTR. Co-transfection significantly decreased luciferase activity (P = 0.043), while mutant constructs failed to do so, confirming target specificity (figure 4C–D).

**Figure 4.**
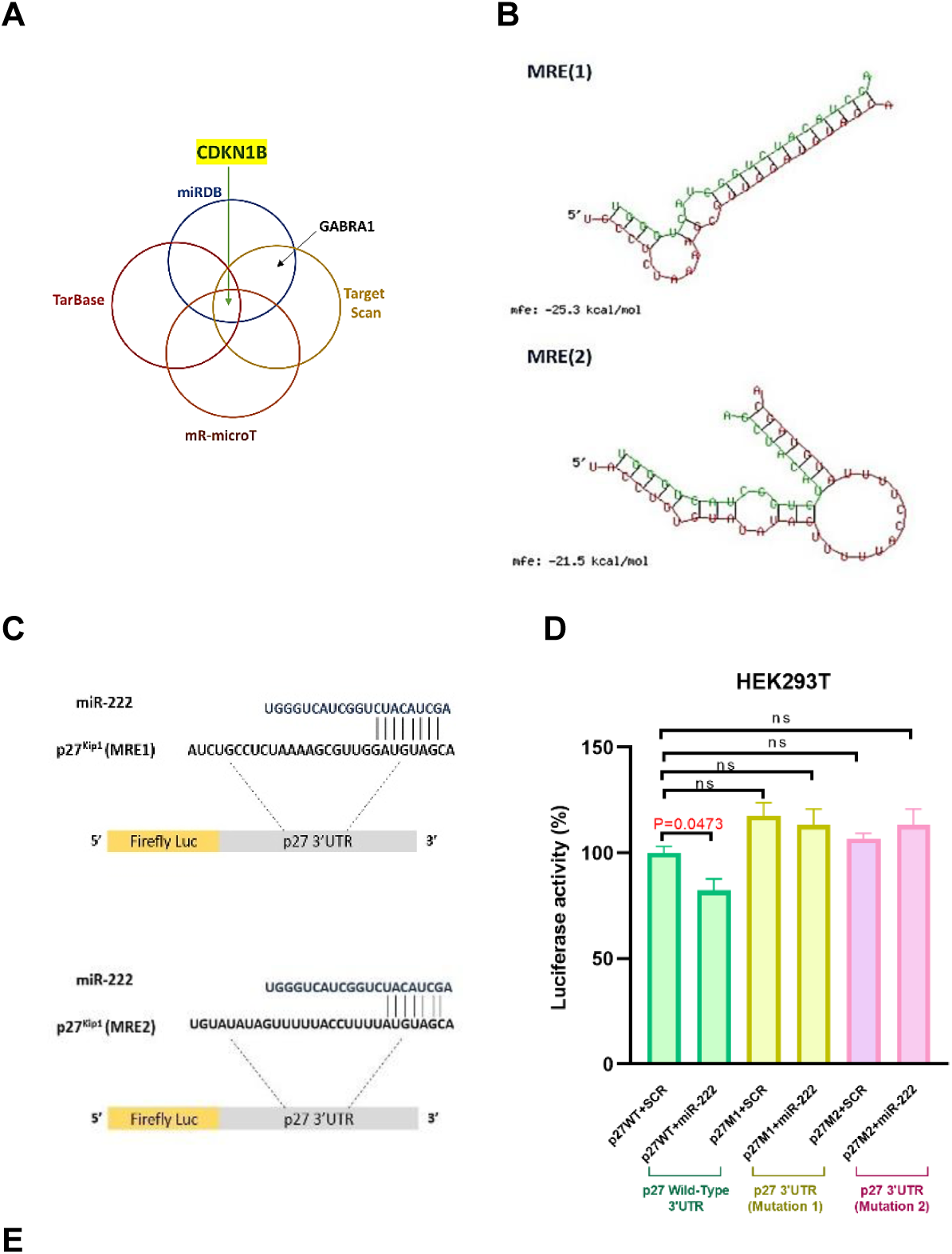
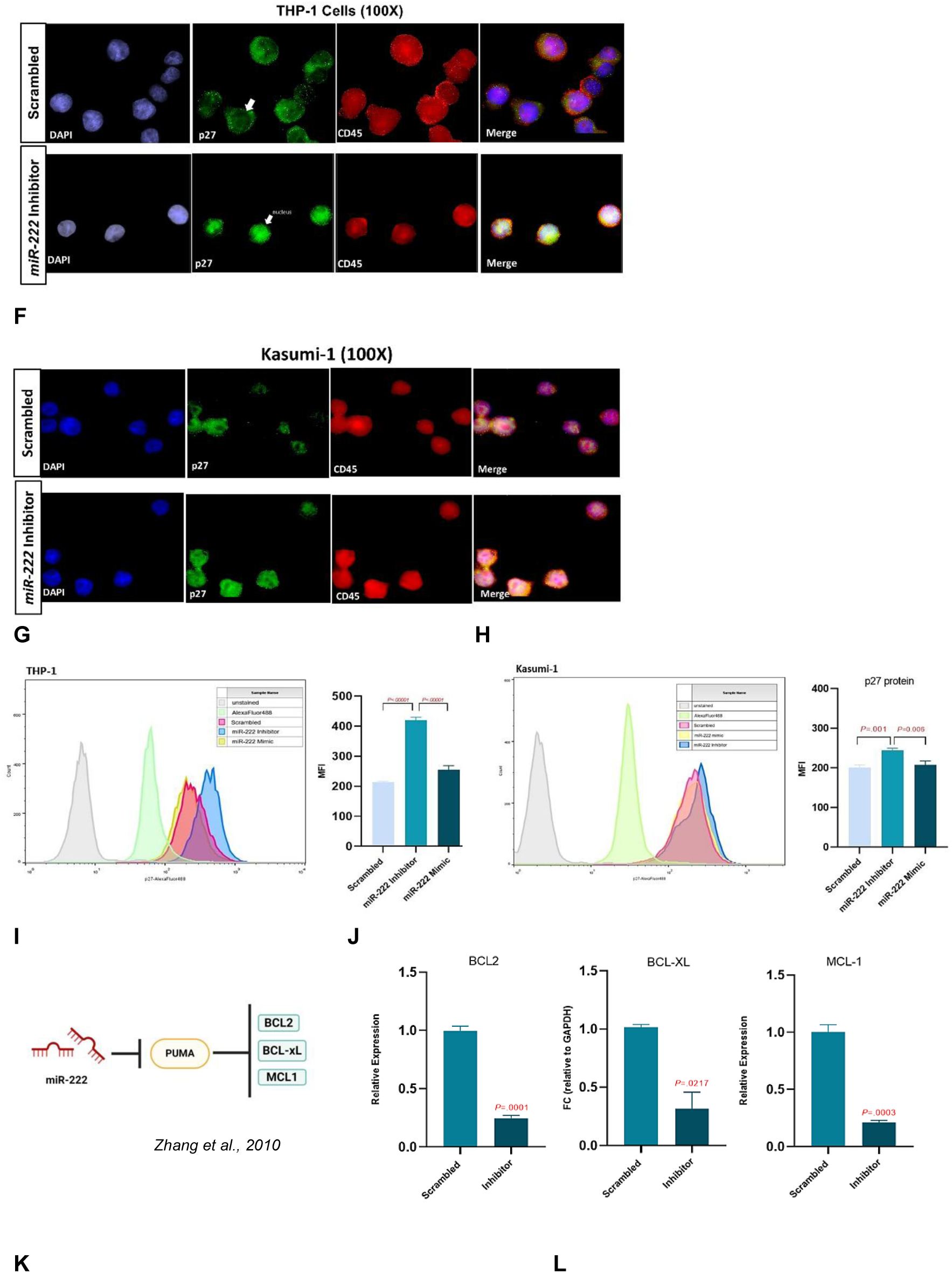
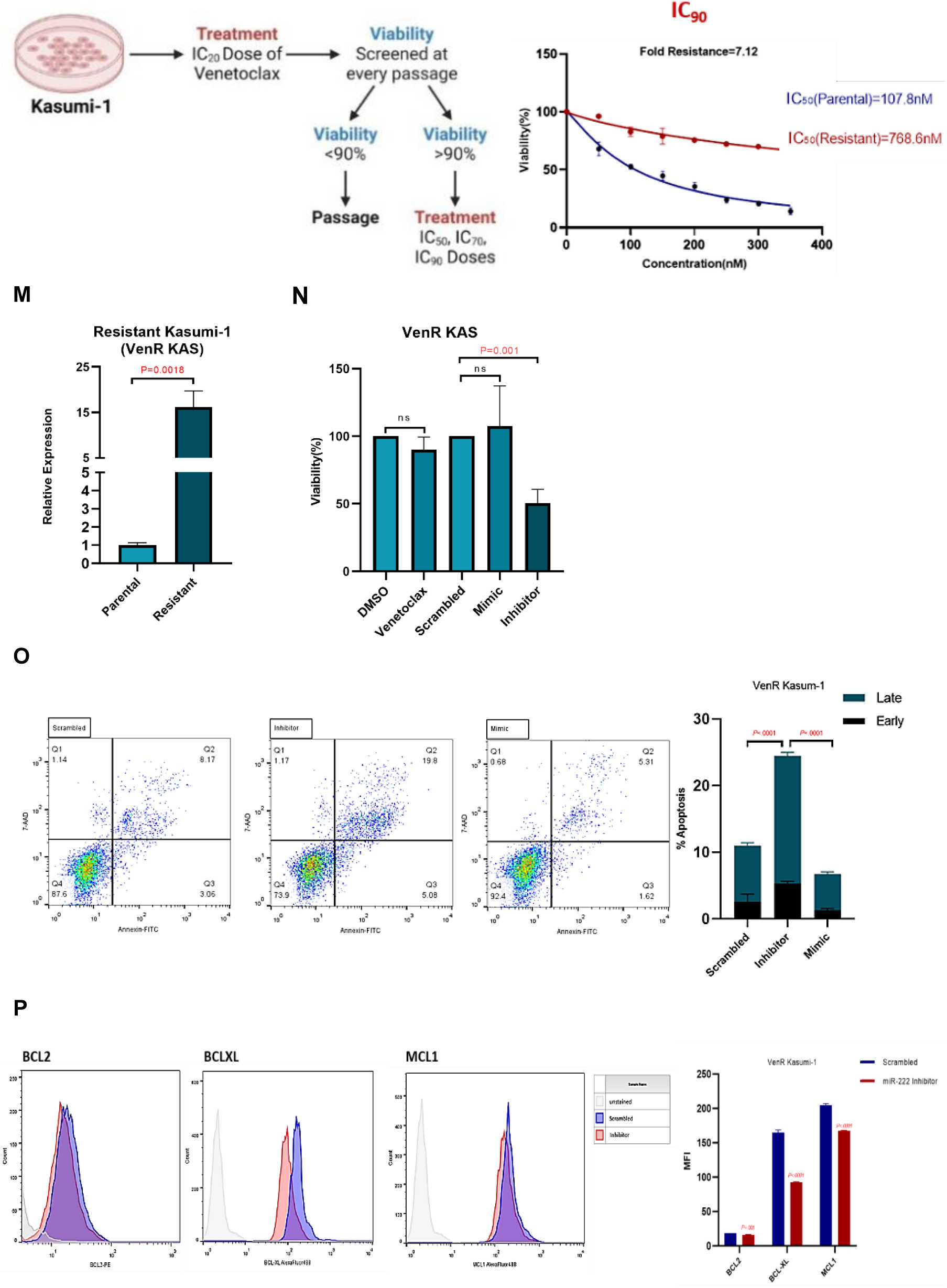
*miR-222* negatively regulates cell cycle inhibitor CKN1B/p27^Kip1^, and promotes venetoclax resistance phenotype in t(8;21) AML Kasumi-1 cell line. A. CDKN1B or p27^Kip1^ was predicted as a 3’UTR target of *miR-222* in 4 different microRNA target prediction databases (miRDB, TargetScan, mR-microT and TarBase). **B.** There are two *miR-222* regulatory elements in the 3’UTR of p27^Kip1^, MRE (1) and MRE (2). RNAHybrid structures showing the putative interaction between MRE1 (8-mer) and MRE2 (7-mer) in the *p27^Kip1^* 3’UTR and *miR-222*. **C.** Depiction of the luciferase assay vector construct using for studying the interaction between *miR-222* and *p27 ^Kip1^*. Both MRE regions of p27 ^Kip1^ were clones 3’ of the firefly luciferase gene in the pmirGLO vector. **D.** Dual Luciferase Assay was performed in HEK293T cells, and the introduction of *p27^Kip1^* 3’UTR containing both MREs along with *miR-222* mimics resulted in a significant decrease in luciferase activity (P=0.043), while *miR-222* mimics, when introduced with mutated MREs did not result in a change of luciferase activity. **E.** Immunofluorescence images depicting that the transfection of *miR-222* inhibitor into THP-1 cells leads to a transport of p27^Kip1^ into the nucleus, from the cytoplasm. **F.** Similarly, increased expression of p27^Kip1^ in the nucleus was observed in Kasumi-1 upon transfection of cells with *miR-222* inhibitor. **G.** Flow cytometry analysis showed that downregulation of *miR-222* in THP-1 cells results in a significant increase in p27^Kip1^ expression (P<0.0001). **H**. Flow cytometric analysis of Kasumi-1 cells after the downregulation of *miR-222* shows increased expression of p27^Kip1^ (P=0.001). **I.** Schematic: *miR-222* induces BCL-2, BCL-xL and MCL-1 expression via post-transcriptional silencing of PUMA. **J.** Downregulation of *miR-222* in Kasumi-1 cells resulted in significant decrease of anti-apoptotic genes BCL-2(P=0.0001), BCL-XL(P=0.0217) and MCL-1(P=0.0003). **K.** Schematic representation of the approach used to develop venetoclax-resistant Kasumi-1 cells **L.** Dose–response curves showing cell viability in parental (blue) and drug-resistant (red) AML cells treated with increasing concentrations of venetoclax. The IC₅₀ of resistant cells (768.6 nM) was significantly higher than that of parental cells (107.8 nM), indicating a ∼7.1-fold increase in resistance. **M.** Resistant Kasumi-1 cells expressed significantly higher levels of *miR-222* (P=0.0018). **N.** The viability of venetoclax resistant Kasumi-1 cells was significantly reduced when treated with *miR-222* inhibitor (P=0.001). **O.** Resistant Kasumi-1 cells in which *miR-222* was inhibited exhibited significantly higher levels of apoptosis (P<0.0001). **P.** The inhibition of *miR-222* in venR Kasumi-1 cells led to a decrease in protein levels of BCL-2 (P=0.001), BCL-(P=0.0001) and MCL-1(P=0.0001). Statistical significance was assessed using unpaired two-tailed Student’s t-test for comparisons between two groups and one-way ANOVA with Tukey’s/Dunnet’s post hoc test for multiple group comparisons. Data are presented as mean ± SD. P-values < 0.05 were considered statistically significant.

To assess whether *miR-222* inhibition alters p27 protein levels, we transfected AML cell lines with *miR-222* inhibitors. Immunofluorescence imaging demonstrated increased nuclear p27 in both THP-1 (figure 4E) and Kasumi-1 cells (figure 4F). Flow cytometry further confirmed increased p27 protein levels in *miR-222* inhibitor-treated THP-1 and Kasumi-1 cells (P = 0.0001 and P = 0.001, respectively) (figure 4G, H), supporting the role of *miR-222* in suppressing p27 expression. In parallel, we investigated whether *miR-222* modulates apoptotic regulators. Upon *miR-222* inhibition, Kasumi-1 cells exhibited significant downregulation of BCL-2 (P = 0.0001), BCL-xL (P = 0.0217), and MCL-1 (P = 0.0003) (figure 4I–J).

Considering the clinical relevance of BCL-2 inhibition, we developed venetoclax-resistant Kasumi-1 cells (ven-R) via stepwise dose escalation (IC20–IC90) (figure 4K). Resistant cells showed a 7.12-fold higher IC50 compared to parental cells (768.6 nM vs. 107.8 nM) (figure 4L; supplementary figure 3D). Transcript analysis of ven-R Kasumi-1 cells revealed significant upregulation of BCL-xL (P = 0.0002) and MCL-1 (P = 0.0002), at transcript (supplementary figure 3E) and protein levels (BCL-xL, P = 0.0033; MCL-1, P < 0.0001) (supplementary figure 4F). Interestingly, *miR-222* expression was significantly elevated in ven-R cells compared to parental cells (P=0.0018) (figure 4M). Functional inhibition of *miR-222* in ven-R Kasumi-1 cells significantly reduced viability (P=0.001) and increased apoptosis (P=0.0001) in venetoclax resistant cells (figure 4N, O). This was accompanied by a marked reduction in BCL-2 (P = 0.001), BCL-xL (P < 0.0001), and MCL-1 (P = 0.0001) protein levels (figure 4P), indicating that *miR-222* facilitates survival under therapeutic stress.

### Epigenetic restoration of *HOTAIRM1* expression sensitizes venetoclax-resistant AML cells to apoptosis

Building on our earlier finding that *HM1V2* negatively regulates *miR-222*, we investigated the significance of the *HOTAIRM1*-*miR-222* regulatory interaction in the context of venetoclax resistance. Baseline *HM1V2* levels were comparable between parental and Ven-R Kasumi-1 cells (figure 5A). However, overexpression of *HM1V2* significantly reduced cell viability (P<0.0001; figure 5B) and increased apoptosis, as indicated by Annexin V/7-AAD staining (P = 0.0022; figure 5C). Mechanistically, *HM1V2* overexpression suppressed anti-apoptotic mediators of the BCL-2 family, including BCL-2 (P = 0.0018), BCL-xL (P = 0.02), and MCL-1 (P = 0.037) at the transcript level (figure 5D), with corresponding reductions in protein levels confirmed by flow cytometry (P < 0.0001 for all; figure 5E). We next examined the expression of key DNA and histone modifying enzymes. The transcript levels of *DNMT3A, DNMT3B, HDAC1,* and *HDAC2* were significantly upregulated in t(8;21) AML compared to controls (figure 5F). We hypothesized that epigenetic repression might underlie *HM1V2* silencing in resistant cells and may be reversible through epigenetic therapy (figure 5H). Accordingly, treatment of Ven-R Kasumi-1 cells with the DNMT inhibitor azacytidine or the pan-HDAC inhibitor panobinostat restored *HOTAIRM1* expression (P = 0.008 and P < 0.0004, respectively), significantly reduced *miR-222* expression levels (P=0.03 and P=0.0001, respectively) (supplementary figure 3H), and induced apoptosis (figure 5J). Combined treatment produced a significantly greater apoptotic response than either agent alone (P < 0.0001), with synergy confirmed by a combination index of 0.64 (figure 5K). These findings demonstrate that restoring *HM1V2* expression not only suppresses leukemic phenotype such as increased proliferation and resistance to apoptosis, but can also significantly affects the viability of venetoclax resistant cells (figure 5L).

**Figure 5.**
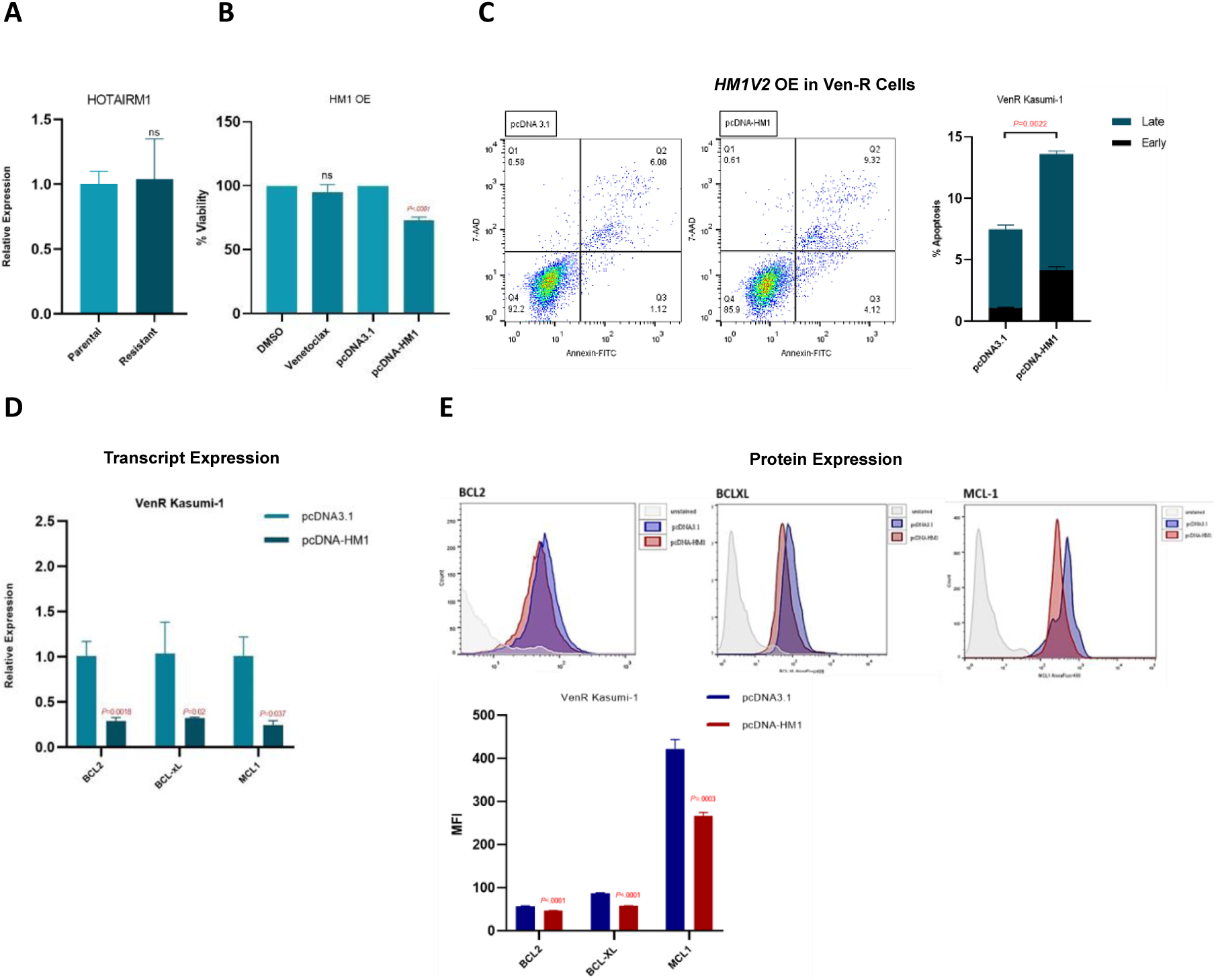
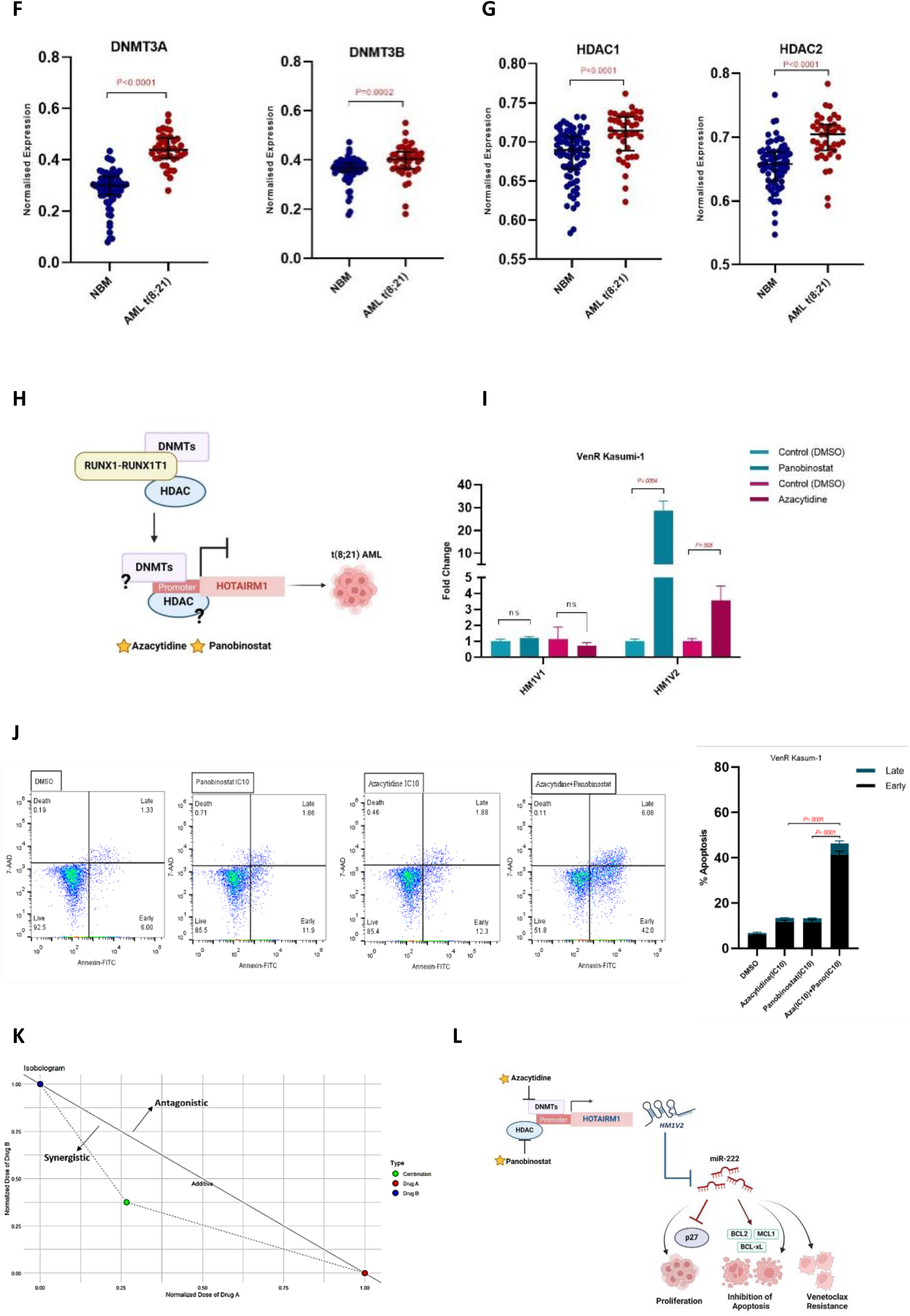
Rescue of *HOTAIRM1* expression abrogates resistance phenotype in Kasumi-1 cells. **A.** The expression of *HM1V2* in venetoclax resistant Kasumi-1 is not different from sensitive Kasumi-1 cells. **B.** *HM1V2* over-expression results in decreased viability of venR Kasumi-1 cells (P<0.0001). **C.** Overexpression of *HM1V2* in Ven-R AML cells leads to significantly increased apoptosis levels (P=0.0022). **D.** A decrease in transcript levels of *BCL-2* (P=0.0018), *BCL-xL* (P=0.02) and *MCL-1* (P=0.03) was observed when *HM1V2* was overexpressed in venR Kasumi-1 cells. **E.** Quantification of protein levels in VenR Kasumi-1 cells after *HM1V2* overexpression indicated significant decrease in i.e.BCL-2 (P<0.0001), BCL-xL(P<0.0001) and MCL-1(P=0.003) **F, G.** *DNMT3A* and *DNMT3B* as wel as *HDAC1* and *HDAC2* are upregulated in t(8;21) AML. **H.** Treatment of cells with Azacytidine (DNMT3 inhibitor) and Panobinostat (HDAC inhibitor) can de-repress silenced gene *HOTAIRM1* **I.** Quantification of *HM1V1 & HM1V2* after treatment of venR Kasumi-1 cells with Azacytidine and Panobinostat. **J.** The combination of Azacytidine and Panobinostat on VenR cells leads to an increased percentage of apoptosis compared to monotreatment (P<0.0001). **K.** Combination Index (CI) of Azacytidine and Panobinostat on VenR cells was found to be 0.64 which indicates a synergestic effect of both drugs in inducing apoptosis. Schematic: *HOTAIRM1*-miR222 pathway and regulation of venetoclax resistance. Statistical significance was assessed using unpaired two-tailed t-tests for pairwise comparisons and combination index (CI) was calculated using the Chou–Talalay method, with CI < 1 indicating synergy. All experiments were performed in biological triplicates, and a P-value < 0.05 was considered statistically significant.

### Expression of *HOTAIRM1-miR-222* and correlation with patient survival outcome

Following *in vitro* functional characterisation of the *HOTAIRM1*–*miR-222* axis and its dysregulation in AML, we evaluated its clinical relevance. We focussed on *miR-222* due to the consistent and significant upregulation across AML and multiple malignancies. The expression of both isoforms of *HOTAIRM1* showed no correlation with patient survival outcome (supplementary table 2). However, Kaplan-Meier survival analysis showed that high *miR-222* expression was significantly associated with inferior event-free survival (EFS; HR, 1.92; 95% CI, 1.219–3.053; P = 0.005) and overall survival (OS; HR, 2.09; 95% CI, 1.290–3.414; P = 0.002) (figure 6A, B), in the current study cohort (i.e. IRCH cohort). These associations were maintained in the t (8;21) AML subgroup, where high *miR-222* was linked to worse EFS (HR, 3.46; 95% CI, 1.431–8.391; P = 0.004) and OS (HR, 2.34; 95% CI, 0.907–6.035; P = 0.03) (figure 6 C, D). In an independent multi-institutional cohort i.e. TARGET-AML, no significant association was observed between *miR-222* and EFS as well as OS in the overall AML population (figure 6E, F). However, among patients with t(8;21) AML, high *miR-222* expression was associated with inferior EFS (HR, 2.51; 95% CI, 1.044–6.429; P = 0.0479), as well as OS (HR, 2.9; 95% CI, 1.032–8.601; P = 0.009 (figure 6 G,H).

**Figure 6.**
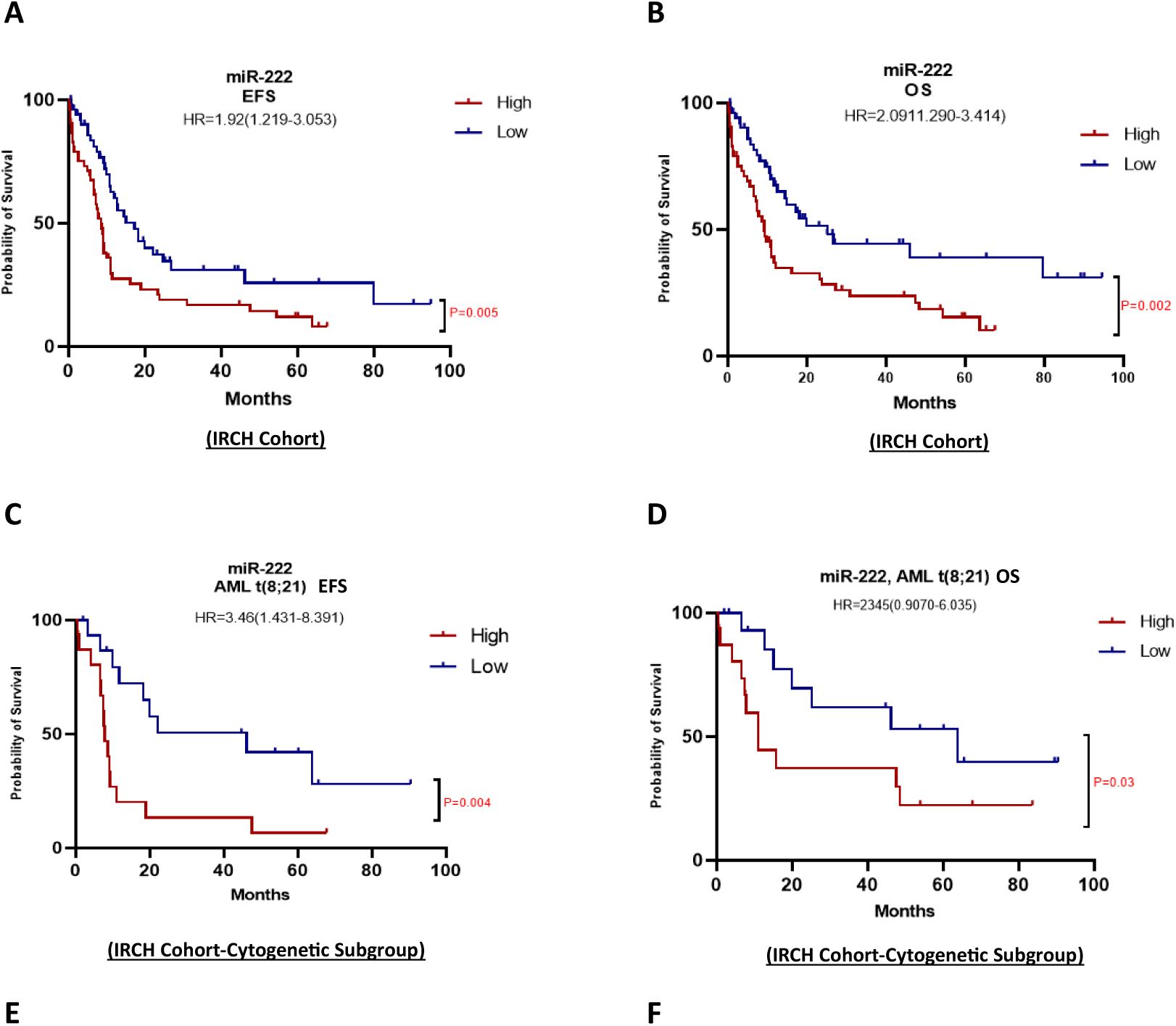
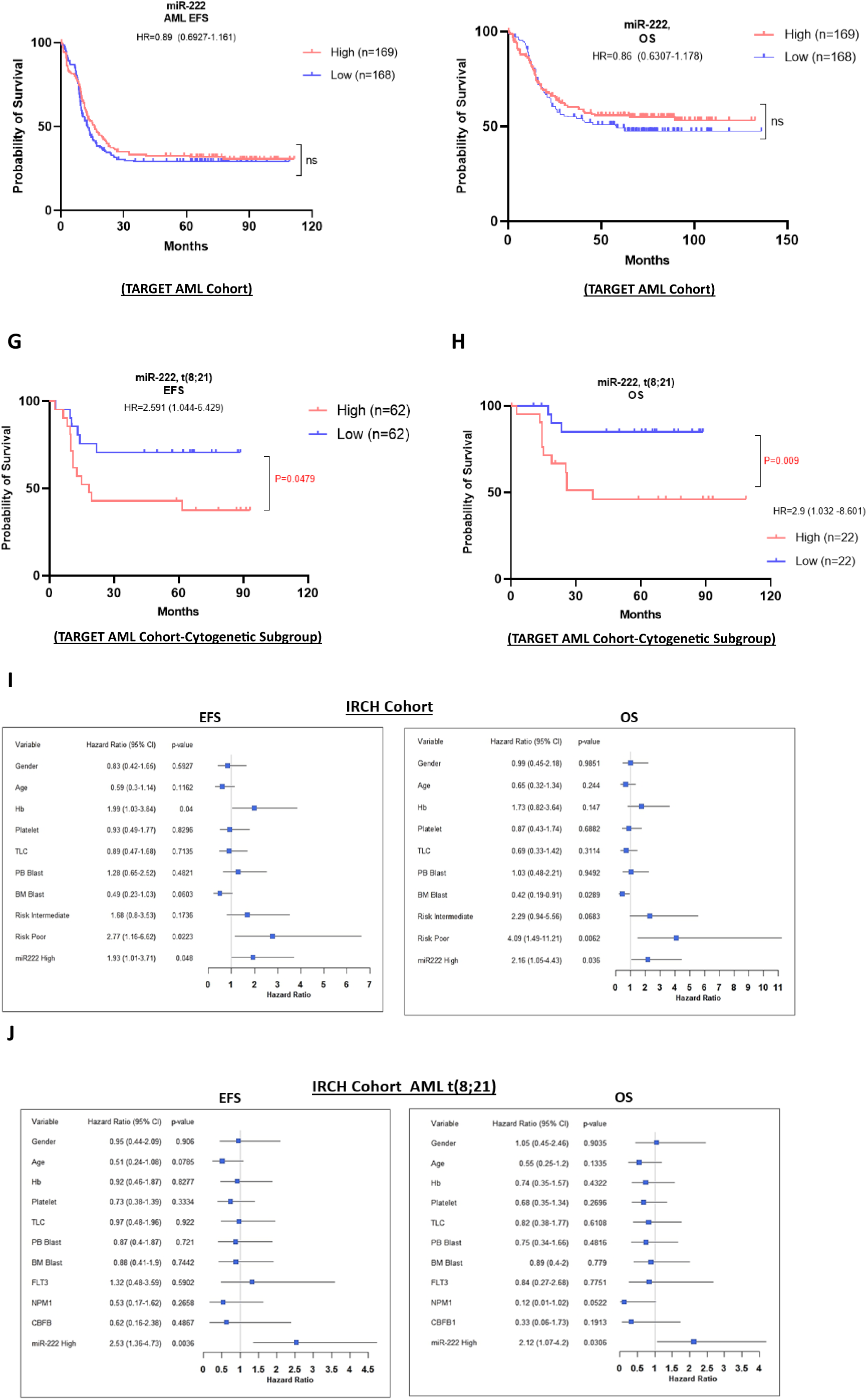
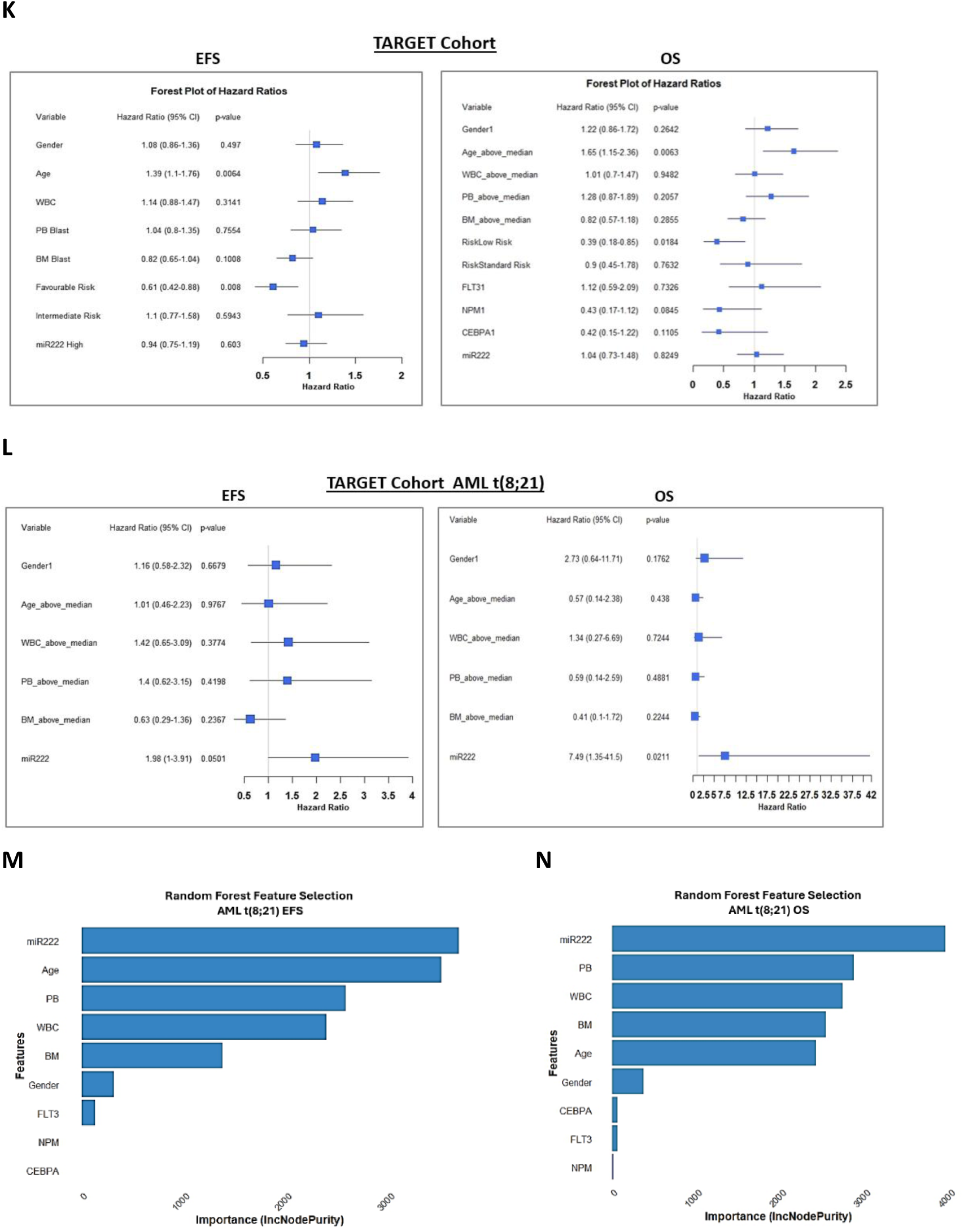
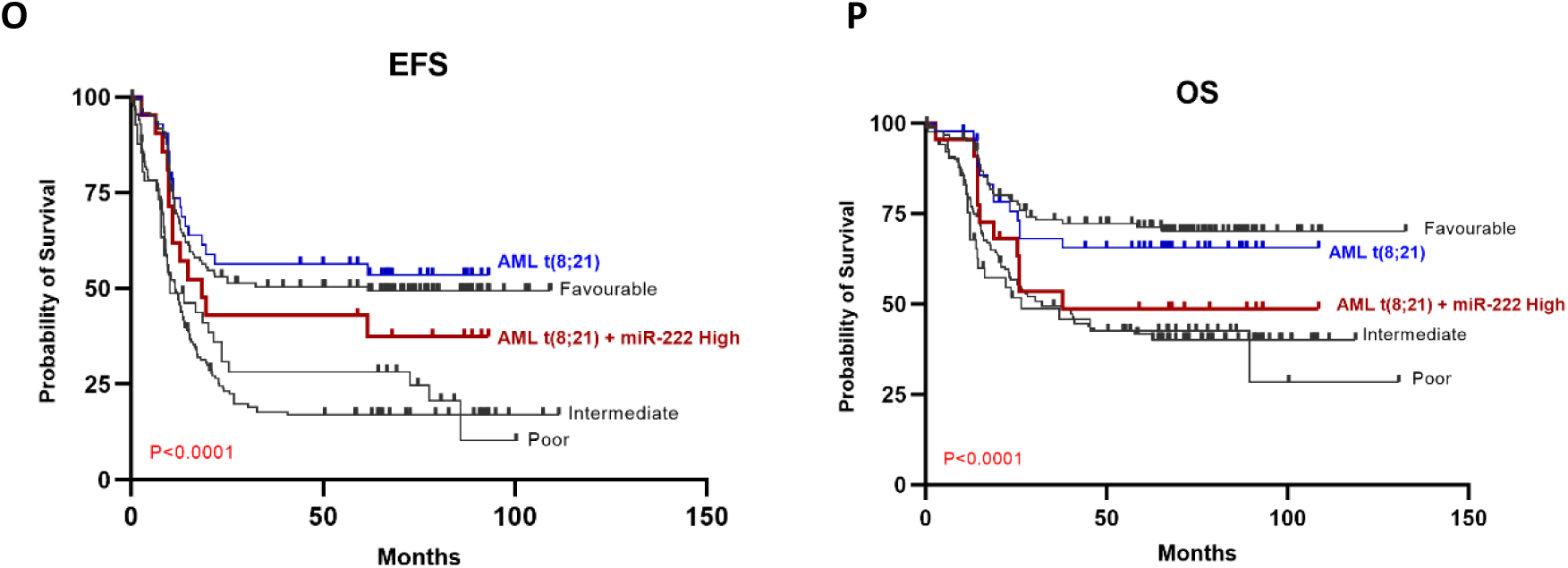
Univariable and multivariable analysis of high versus low *miR-222* expression on survival outcomes in two cohorts of AML patients. **A,B,C,D.** Event free survival and overall survival of patients stratified by *miR-222* expression in all AML patients as well as the t(8;21) AML subgroup in the IRCH cohort, where high *miR-222* expression is associated with poor overall survival (P<0.01). **E,F,G,H.** EFS and OS of patients with AML from the entire TARGET AML cohort as well from the t(8;21) AML subtype. High *miR-222* expression and its association with poor EFS and OS in the t(8;21) AML subtype was confirmed (P<0.05). **I,J,K,L**. Forest plots for hazard ratios of high versus low *miR-222* expression in overall as well as t(8;21) AML cohorts, demonstrating its independent association with survival outcomes in the IRCH cohort as well as in the TARGET cohort. **M, N**. Random Forest Feature Selection for predicting EFS and OS in the t(8;21) AML cohort. **M.** shows the importance of features like *miR-222*, age, PB, WBC, and gender for EFS prediction, while **N.** highlights *miR-222*, PB, WBC, BMI, and age as the most influential factors in predicting OS, with *miR-222* as a key prognostic factor in t(8;21) AML. **O, P.** Kaplan-Meier curves depicting EFS and OS showing that AML t(8;21) patients with high *miR-222* expression have lower survival probabilities compared to all AML t(8;21) patients. Statistical analyses included Kaplan– Meier survival estimates with log-rank tests for group comparisons (P<0.05 considered significant) and Cox proportional hazards models for univariable and multivariable analysis. Random Forest algorithm was used for feature selection, and variable importance was ranked based on mean decrease in Gini impurity. Analyses were performed separately for overall AML and the t(8;21) AML subtype across both IRCH and TARGET cohorts.

Multivariable Cox regression analysis confirmed that high *miR-222* expression was an independent predictor of poor EFS (HR, 1.31; 95% CI, 1.013–1.717; P = 0.048) and OS (HR, 2.16; 95% CI, 1.054–4.43; P = 0.036) in the current study cohort (figure 6I), after adjusting for age, haemoglobin, gender, blast percentage and cytogenetic risk, among other clinical characteristics. In the t(8;21) AML subgroup, high *miR-222* expression also retained independent prognostic value, significantly predicting worse EFS (HR, 2.53; 95% CI, 1.364–7.3; P = 0.036) and OS (HR, 2.12; 95% CI, 1.074–4.2; P = 0.0306) (figure 6J).

In the TARGET cohort *miR-222* showed a borderline association with worse EFS (HR, 1.98; 95% CI, 1.331–2.99; P = 0.051) but was significantly associated with OS in t(8;21) AML (HR, 7.49; 95% CI, 1.345–41.5; P = .0211) (figure 6L).

Random forest based feature selection ranked *miR-222* as the top prognostic feature for both EFS and OS in t(8;21) AML, outperforming clinical variables such as age, peripheral blood blast count, and white blood cell count (figure 6M, N).

Patients with AML t(8;21) and high miR-222 expression exhibited survival outcomes comparable to those in intermediate- and poor-risk groups, suggesting that elevated miR-222 may offset the favorable prognosis typically conferred by the t(8;21) cytogenetic subtype (Figure 6O, P; P<0.001) (Table 2).

**Table 2.**
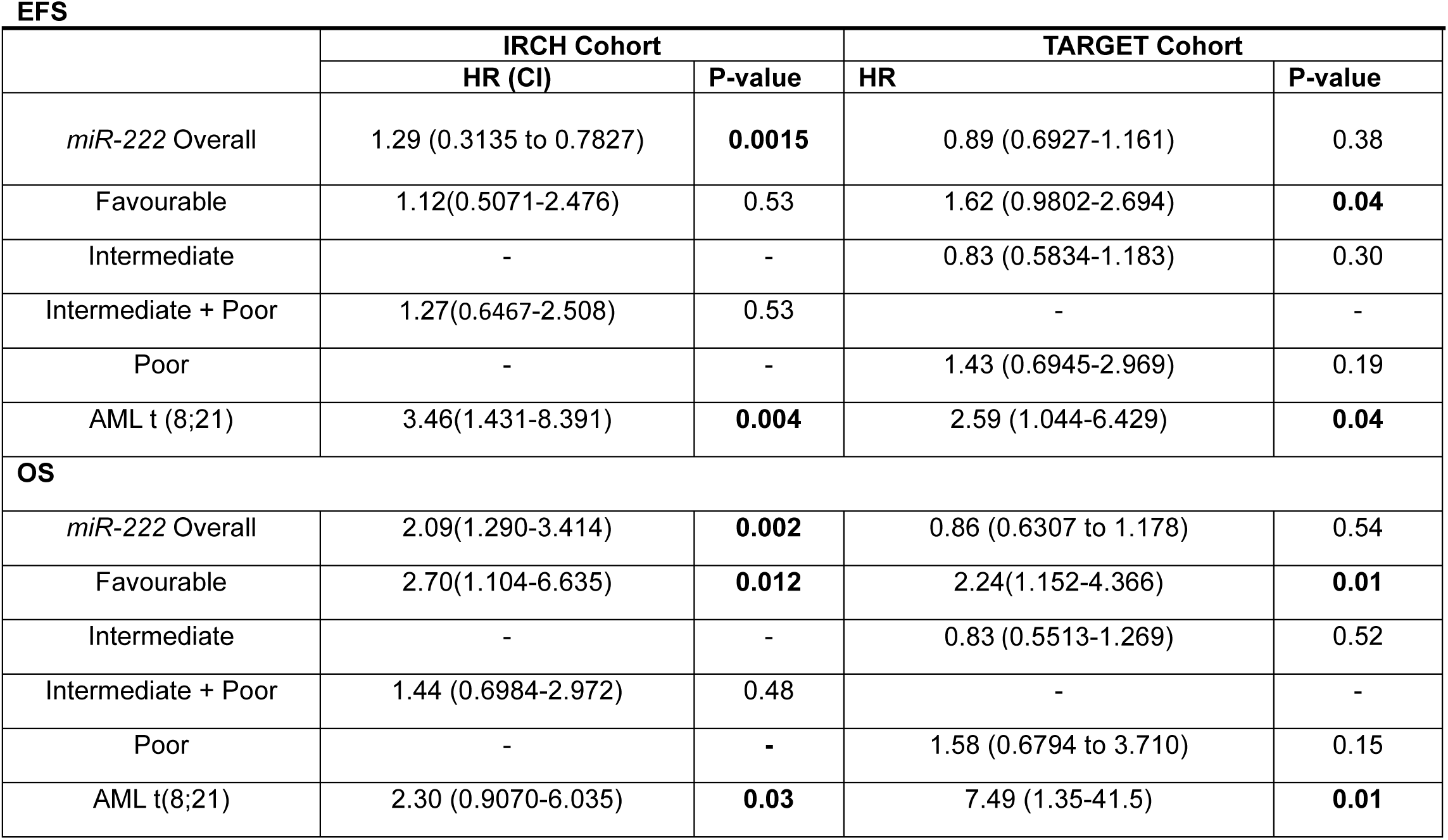
Comparison of *miR-222* expression and survival outcome (internal vs. external cohort)

|  | IRCH Cohort |  | TARGET Cohort |  |
| --- | --- | --- | --- | --- |
|  | HR (CI) | P-value | HR | P-value |
| *miR*-222 Overall | 1.29 (0.3135 to 0.7827) | **0.0015** | 0.89 (0.6927-1.161) | 0.38 |
| Favourable | 1.12(0.5071-2.476) | 0.53 | 1.62 (0.9802-2.694) | **0.04** |
| Intermediate | - | - | 0.83 (0.5834-1.183) | 0.30 |
| Intermediate + Poor | 1.27(0.6467-2.508) | 0.53 | - | - |
| Poor | - | - | 1.43 (0.6945-2.969) | 0.19 |
| AML t (8;21) | 3.46(1.431-8.391) | **0.004** | 2.59 (1.044-6.429) | **0.04** |
| OS |  |  |  |  |
| *miR*-222 Overall | 2.09(1.290-3.414) | **0.002** | 0.86 (0.6307 to 1.178) | 0.54 |
| Favourable | 2.70(1.104-6.635) | **0.012** | 2.24(1.152-4.366) | **0.01** |
| Intermediate | - | - | 0.83 (0.5513-1.269) | 0.52 |
| Intermediate + Poor | 1.44 (0.6984-2.972) | 0.48 | - | - |
| Poor | - | - | 1.58 (0.6794 to 3.710) | 0.15 |
| AML t(8;21) | 2.30 (0.9070-6.035) | **0.03** | 7.49 (1.35-41.5) | **0.01** |

## Discussion

Although the t(8;21) translocation is traditionally considered a favorable-risk abnormality in AML, accumulating evidence indicates that this subtype is clinically heterogeneous, with a subset of patients experiencing poor outcomes independent of cytogenetic aberrations^42,43^. Dysregulation of hematopoietic lineage specific lncRNAs is increasingly recognized as a contributor to leukemic progression^20^.

In this study, we identified significant downregulation of two myeloid differentiation associated lncRNAs, *HOTAIRM1* and *LINC00173*, in t(8;21) AML. We chose to focus on *HOTAIRM1* and dissect its functional relevance in this leukemic subtype because it is transcriptionally regulated through direct binding of the myeloid transcription factor PU.1 to its promoter, and PU.1 itself is repressed by the RUNX1–RUNX1T1 oncofusion protein ^24,44^. Supporting this, our *in silico* analysis across hematopoietic lineages revealed that both *HOTAIRM1* and its regulator PU.1 exhibit peak expression in mature myeloid populations, but are downregulated in t(8;21) AML.

*HOTAIRM1* has a short isoform, *HOTAIRM1 variant 2* (*HM1V2*) with a role in myeloid differentiation and a long isoform *HOTAIRM1 variant 1* (*HM1V1*) associated with neuronal differentiation (Figure 1J) ^24,45^. Aberrant levels of RNA isoforms are known to play a specific and central regulatory role in AML drug resistance^46^. In our study, we focused on the lncRNA isoform *HM1V2,* and demonstrated that the RUNX1-RUNX1T1-PU.1-*HM1V2* axis forms a mechanistic link between the oncofusion protein and impaired myeloid maturation through *HM1V2* dysregulation.

Building on this observation, and on our experimental data which showed *HM1V2* upregulation following epigenetic drug treatment, we propose that in t(8;21) AML, *HM1V2* is transcriptionally silenced through a dual mechanism: repression of PU.1 by the RUNX1-RUNX1T1 fusion protein, and epigenetic alterations involving deregulated histone deacetylase and DNA methyltransferase activity, both mechanisms known to be associated with the t(8;21) translocation ^47^.

The restoration of *HM1V2* expression in t(8;21)-positive Kasumi-1 cell line, led to accumulation of cells in the G0/G1 phase of the cell cycle and induction of apoptosis, highlighting a tumor suppressive role. Additionally, the lncRNA *HM1V2* negatively regulates oncogenic microRNA *miR-222*, establishing a link between disrupted differentiation pathways and aberrant survival signalling in t(8;21) AML.

Importantly, *miR-222* is a well-characterized oncomiR, upregulated in multiple solid tumors where it promotes tumor progression, therapy resistance, and poor clinical outcomes.^48–50^. Its prognostic relevance has been validated by meta-analyses, supporting its utility as both a biomarker and a therapeutic target in cancer ^35,35,51^. Extending these observations, our expression analysis across various solid tumor cell lines, including triple-negative breast cancer (TNBC), estrogen receptor–positive breast cancer, thyroid cancer, ovarian cancer, and B-ALL showed consistently elevated levels of *miR-222* compared to normal cells (Supplementary Figure 2E). This was further supported by TCGA data, where miR-222 was upregulated in 16 out of 22 analyzed solid tumor types (supplementary figure 2F), reinforcing its broad oncogenic significance.

In addition to this, we found that *miR-222* is detectable at significantly higher levels in plasma-derived exosomes from patients with AML compared to healthy individuals, suggesting its potential utility as a minimally invasive biomarker. This finding highlights the utility of exosomal *miR-222* as a valuable tool for non-invasive disease monitoring^52,53^.

Our *in vitro* studies confirm that *miR-222* binds the 3′UTR of the cyclin-dependent kinase inhibitor i.e. *p27*, promoting cell survival in AML through post-transcriptional silencing of p27 while sustaining BCL-2 family mediated anti-apoptotic signalling. In venetoclax-resistant AML cells, elevated *miR-222* levels maintain BCL-xL and MCL-1 expression. Inhibiting *miR-222* restores p27 expression, reduces BCL-2 family protein levels, and re-sensitizes venetoclax resistant cells to apoptosis.

In line with our findings, Zhang et al. reported that *miR-222* promotes AML cell proliferation and inhibits apoptosis. They identified the targeting of Axin2 and activation of Wnt/β-catenin pathway via *miR-222*, as an additional mechanism through which *miR-222* enhances leukemic cell survival^49^.

Mutations in the *KIT* gene, which frequently co-occur with t(8;21) AML, have been implicated in promoting venetoclax resistance ^12,54^. Our analysis of the TARGET-AML dataset revealed that four out of fifty-nine patients with t(8;21) AML harboured KIT mutations, all of which exhibited high (above-median) miR-222 expression levels. Additionally, the Kasumi-1 cell line used in our experiments, which models t(8;21) AML, also carries a *KIT* mutation, further supporting a possible interplay between *KIT* mutation status and *miR-222* mediated resistance mechanisms^53^.

We hypothesised that the restoration of *HM1V2* expression could overcome *miR-222* mediated venetoclax resistance phenotype. Treatment of resistant cells with the DNA hypomethylating agent azacytidine in combination with the HDAC inhibitor panobinostat restored *HM1V2* expression and sensitised resistant cells to apoptosis, supporting a synergistic epigenetic approach to address resistance. This is particularly relevant as venetoclax is increasingly used in frontline AML treatment, with concerns about emerging resistance, particularly in the t(8;21) AML subtype ^9,11,16^.

To assess the prognostic impact of the *HOTAIRM1*–*miR-222* axis in t(8;21) AML, we examined whether expression levels were associated with clinical outcomes. While *HM1V2* expression showed no significant correlation with survival (supplementary table 3), elevated *miR-222* levels independently predicted both poorer event-free and overall survival in our institutional pediatric cohort and were validated in the multi-institutional TARGET-pediatric AML dataset. Machine learning-based feature selection ranked *miR-222* as the top prognostic variable within this cytogenetic subgroup, further supporting its utility for refined risk stratification. Although contemporary prognostic models in AML integrate cytogenetics, gene mutations, and clinical parameters, no non-coding RNA expression signature has to date demonstrated independent prognostic value specifically within the t(8;21) subgroup.

This study has certain limitations. Our institutional pediatric AML cohort comprised a relatively small sample size of 106 patients, which could limit the generalizability of the results. To address this and improve the robustness of our observations, we validated the prognostic significance of *miR-222* in the Therapeutically Applicable Research to Generate Effective Treatments (TARGET) AML dataset, a large, multi-institutional cohort of pediatric AML patients with comprehensive genomic and clinical annotation. Larger, prospective studies will be necessary to confirm these findings in diverse populations. Additionally, the development of an internationally standardized assay to accurately quantify *miR-222* in clinical samples will be essential before it can be incorporated into routine risk stratification in AML.

Another important consideration is that the role of the bone marrow microenvironment, which is known to confer protective niches for leukemic cells, was not addressed in this study. Interactions between leukemic blasts and stromal, endothelial, and immune components of the marrow niche promote survival, quiescence, and therapy resistance including resistance to venetoclax through soluble factors, adhesion molecules, and hypoxia-driven signalling^55,56^.

In addition to microenvironmental influences, other cell-intrinsic mechanisms contributing to venetoclax resistance have also been reported. These include acquired mutations in BCL-2 that reduce venetoclax binding affinity, as well as alterations in mitochondrial dynamics and metabolic reprogramming^57,58^. These intrinsic and niche-mediated effects may act in parallel or synergistically with the *HOTAIRM1–miR-222* axis and may represent alternate resistance pathways not captured in our *in vitro* experiments.

Future studies incorporating co-culture systems, patient-derived xenografts, or spatial transcriptomics will be necessary to elucidate how the microenvironment influences non-coding RNA mediated resistance and to explore additional mechanisms contributing to the resistant phenotype.

Overall, our findings indicate that the molecular landscape of t(8;21) AML may be more heterogeneous than previously understood, with non-coding RNAs playing a crucial role in the pathobiology of this subtype. This study highlights the *HOTAIRM1*–*miR-222* axis as an important regulator of venetoclax resistance, which is targetable using epigenetic therapy, and provides strong rationale for integrating non-coding RNA profiling into standard AML risk stratification.

## FUNDING AND ACKNOWLEDGEMENTS

This study was supported by the Young Investigator Award from the Department of Science and Technology - Science and Engineering Research Board (DST-SERB), Government of India; the intramural research grant (IRG) from All India Institute of Medical Sciences, New Delhi, India; and a junior and senior research fellowship from the Council of Scientific and Industrial Research (CSIR), Government of India. We thank Dr. Theodum Debraj, Department of Medical Oncology, AIIMS, New Delhi, India for providing the cell lines used for the microRNA expression analysis. Figures were created using BioRender.com (License: https://BioRender.com/1ma8c77; Wilson, C., 2025).

