## Supplementary Figures and Tables for "The *HOTAIRM1-miR-222* Axis Regulates Venetoclax Resistance and Defines a High-Risk Subset in Pediatric t(8;21) Acute Myeloid Leukemia"

**Supplementary Methods:**

1. **Sample Collection and Processing**

Peripheral blood (PB) samples were collected from AML patients and healthy controls in EDTA-coated vials (DBO Phlecocuum®, Indore, India) and processed immediately upon receipt. Red blood cell lysis was performed using RBC lysis buffer (155 mM NH4Cl, 12 mM NaHCO3, 0.1 mM EDTA). The samples were incubated in lysis buffer for 15 minutes, centrifuged at 1,500 rpm for 10 minutes and the pellet was washed twice with 1X phosphate-buffered saline (PBS, Gibco®, MA, USA) to remove any residual lysed red blood cells.

For RNA isolation, the pellet was resuspended in 1 ml TRIzol® (Thermo Fisher Scientific, MA, USA) and processed according to the manufacturer's protocol. RNA concentration and purity were assessed using NanoDrop 2000 (Thermo Fisher Scientific, MA, USA).

2. **RNA Extraction and Quantitative PCR (qPCR)**

Total RNA was extracted using TRIzol® Reagent (Thermo Fisher Scientific, MA, USA) following the manufacturer’s protocol. RNA integrity was confirmed using a NanoDrop spectrophotometer (Thermo Fisher Scientific, Wilmington, DE, USA). For mRNA analysis, cDNA synthesis was performed with the High-Capacity cDNA Reverse Transcription Kit (Applied Biosystems, CA, USA) using 1 µg of RNA in a 20 µl reaction following the manufacturer’s protocol. For microRNAs, cDNA was synthesized using the miScript II RT Kit (Qiagen, Hilden, Germany) using 1 µg of RNA and miScript HiFlex Buffer, followed by incubation at 37°C for 60 minutes and enzyme inactivation at 95°C for 5 minutes. Quantitative real-time PCR (qPCR) for mRNAs and lncRNAs was performed using PowerUp™ SYBR™ Green Master Mix (Thermo Fisher Scientific, MA, USA). The thermal cycling conditions consisted of an initial denaturation at 95°C for 10 minutes, followed by 40 cycles of 95°C for 15 seconds, 55°C for 30 seconds, and 72°C for 30 seconds. A melt curve analysis was performed, incrementally increasing the temperature from 55°C to 95°C, to confirm the specificity of the amplification and the absence of non-specific products. The qPCR reaction for microRNAs was performed using the miScript SYBR Green PCR Kit (Qiagen, Germany) on the LightCycler 480 system (Roche, Basel, Switzerland). The thermal cycling conditions consisted of an initial denaturation at 95°C for 15 minutes, followed by 40 cycles of 95°C for 15 seconds, 55°C for 30 seconds, and 72°C for 30 seconds. The primer sequences used in qPCR have been described in supplementary table 1.

Relative expression levels were calculated using the 2^−ΔΔCt^ method[53] with GAPDH as the endogenous control for mRNA and lncRNA normalization, and U6 as the endogenous control for microRNA normalization. All reactions were performed in triplicate.

3. **Cell Culture**

The AML cell lines Kasumi-1, THP-1, MOLM-13, MOLM-14, HL-60, and KG-1, as well as the non-leukemic control cell line HEK293T, were used in this study. Kasumi-1, was obtained as a gift from collaborator Dr. Sameer Bakhshi, THP-1, MOLM-13 and MOLM-14 were obtained as a gift from collaborator Dr. Sampa Ghose, HL-60 and KG-1 cell lines were obtained from the National Centre for Cell Science (NCCS, Pune, India). HEK293T cells were obtained from collaborator Dr. Jayanth K. Palanichamy.

Cells were maintained under standard culture conditions at 37°C in a humidified incubator with 5% CO2. Kasumi-1 was cultured in RPMI-1640 medium (Gibco, Thermo Fisher Scientific) supplemented with 20% fetal bovine serum (FBS, Gibco) and 1% penicillin-streptomycin (100 U/ml and 100 µg/ml, respectively; Thermo Fisher Scientific, MA, USA). THP-1, MOLM-13, MOLM-14, HL-60 and KG-1 cells were cultured in RPMI-1640 supplemented with 10% FBS, and 1% penicillin-streptomycin. HEK293T cells were cultured overnight in Dulbecco’s Modified Eagle Medium (DMEM) supplemented with 10% fetal bovine serum (FBS) and 1% penicillin-streptomycin. Cell viability was routinely assessed using trypan blue exclusion, and cultures were tested regularly to confirm the absence of mycoplasma contamination.

4. **Immunofluorescence**

Immunofluorescence analysis was performed to assess the expression of p27 in AML cell lines. Following transfection with *miR-222* inhibitors or scrambled controls for 48 hours, Kasumi-1 or THP-1 cells were collected and fixed with 4% paraformaldehyde for 15 minutes at room temperature.

Fixed cells were permeabilized with 0.1% Triton X-100 in phosphate-buffered saline (PBS) for 10 minutes and blocked with 5% bovine serum albumin (BSA) in PBS for 1 hour to prevent nonspecific binding. Cells were incubated with a primary antibody against p27 (1:200 dilution; Affinity Biosciences, Ohio, USA), BCL-2 (BD Biosciences, CA, USA), BCL-xL or MCL-1(Elabscience, Hubei, China) for 1 hour. After washing with PBS, the cells were incubated with Alexa Fluor 488-conjugated secondary antibody (Abcam, Cambridge, UK) for 1 hour at room temperature in the dark. Following secondary antibody incubation, the cells were washed with PBS and counterstained with DAPI (4′,6-diamidino-2-phenylindole) to visualize nuclei. For co-localization studies, CD45-PE (1:100 dilution; BD Biosciences, CA, USA) was added as a cell surface marker to distinguish leukemic cells. Images were acquired using a fluorescence microscope (Leica Microsystems, Wetzlar, Germany) with appropriate filters for DAPI, Alexa Fluor 488, and PE. Image analysis and overlay were performed using ImageJ (NIH, Bethesda, MD, USA).

5. **MTT Assay and IC_50_ determination**

To assess the effects of drug treatments on cell viability, Kasumi-1 cells were seeded in 96-well plates at a density of 1 × 10^5^ cells per well and treated with increasing concentrations of venetoclax (MedChemExpress, NJ, USA) dissolved in DMSO (Missouri, USA). Following 48 hours of treatment, cell viability was determined using the MTT assay (Thermo Fisher Scientific, Waltham, MA, USA). Briefly, 10 μl of MTT reagent (5 mg/ml) was added to each well and incubated for 4 hours at 37°C. Formazan crystals were solubilized in 100μl of dimethyl sulfoxide (DMSO), and absorbance was measured at 570 nm using a microplate reader (BioTek, Vermont, USA). The half-maximal inhibitory concentration (IC50) values were calculated using non-linear regression analysis in GraphPad Prism (v9.0). The drugs Azacytidine (MedChemExpress, NJ, USA), and Panobinostat (MedChemExpress, NJ, USA), were also used for toxicity assays after dissolving in DMSO.

6. **Cloning**

To investigate whether the downregulation of *HOTAIRM1* has significance in the pathobiology of AML, we designed primers to amplify the full-length sequence of *HOTAIRM1* isoforms, *HM1V1* and *HM1V2* (Figure 2A). A band of approximately 700bp was amplified and ligated into the pcDNA3.1+ vector (Supplementary Figure 1A). The clones obtained after ligation were digested using EcoRI and XhoI to confirm presence of the cloned sequence (Supplementary Figure 1B, lane 3). Sanger sequencing demonstrated a 100% similarity of the cloned sequence to the NCBI reference sequence of NR_038367.1 (Supplementary Figure 1C).

The secondary structures of the cloned *HOTAIRM1* sequence as well as the reference NCBI sequence were generated using RNAFold and found to be have similar free energy and stem loop structures (Figure 2B), indicating that the cloned *HOTAIRM1* transcript was similar in secondary structure to the reference transcript.

To demonstrate transfection efficiency, expression analysis of *HOTAIRM1* was performed in AML cell lines using qPCR.

Additionally, flow cytometry analysis was performed to quantify the expression protein levels of CD11b, a known downstream target of *HOTAIRM1*.

**Supplementary Tables**

**Table 1. Primers used in qPCR**

|  | **Gene** | **Type** | **Sequence** |
| --- | --- | --- | --- |
| 1 | *HOTAIRM1* variant 1 *(HM1V1)* | LncRNA | FP:CATCGCGTTGTCATTGGAAC RP: GGGTTCAGGCAAAACAGAC |
| 2 | *HOTAIRM1* variant 2 *(HM1V2)* | LncRNA | FP:CATCGCGTTGTCATTGGAAC RP: CACCCACATTTCAACCCC |
| 3 | *GAPDH* | House-keeping  (mRNA) | FP:CCCAGCAAGAGCACAAGAGG  RP:TGGTACATGACAAGGTGCGG |
| 4 | *BCL-2* | mRNA | FP: GTGGCCTTCTTTGAGTTCGG  RP:GGCCGTACAGTTCCACAAAG |
| 5 | *BCL-xL* | mRNA | FP:GCCACTTACCTGAATGACCACC  RP:GAACCAGCGGTTGAAGCGTTC |
| 6 | *MCL-1* | mRNA | FP:CCAAGAAAGCTGCATCGAACC RP:CAGCACATTCCTGATGCCAC |
| 7 | *CDKN1B  (p27/Kip1)* | mRNA | FP: GCGCAGGAATAAGGAAGCGAC  RP: CGTCTGCTCCACAGAACCG |
| 8 | *hsa-miR222-3p* | microRNA | FP: GACGAGCTACATCTGGCTACTG RP: Universal Primer (Kit) |
| 9 | *hsa-miR148b-3p* | microRNA | FP: GACGTCAGTGCATCACAGAACT  RP: Universal Primer (Kit) |
| 10 | *hsa-mir424-5p* | microRNA | FP: GCAGCAGCAATTCATGTTTTG  RP: Universal Primer (Kit) |
| 11 | *U6* | House-keeping (small RNA) | FP:GCTTCGGCAGCACATATACTAAAAT  RP:GAACGCTTCACGAATTTGCGTG |

**Table 2. Dataset References for Gene Expression Analysis.**

|  | **Accession Number/ Name** | **Used for quantification**  **of gene:** | **Data Type** | **Link** |
| --- | --- | --- | --- | --- |
| 1. | GSE98791 | *HOTAIRM1, PU.1* | Micro-array | https://www.ncbi.nlm.nih.gov/geo/query/acc.cgi?acc=GSE98791 |
| 2. | GSE116256 | *HOTAIRM1* | Single-Cell RNA-sequencing | https://www.ncbi.nlm.nih.gov/geo/query/acc.cgi |
| 3. | GSE13159 (Leukemia MILE) | *HOTAIRM1, PU.1* | Micro-array | https://www.ncbi.nlm.nih.gov/geo/query/acc.cgi?acc=GSE13159 |
| 4. | GSE117090 | *miR-221, miR-221* | Micro-array | https://www.ncbi.nlm.nih.gov/geo/query/acc.cgi?acc=GSE117090 |
| 5. | GSE51908 | *miR-221, miR-222* | Micro-array | https://www.ncbi.nlm.nih.gov/geo/query/acc.cgi |
| 6. | TARGET-AML | *miR-221, miR-222* | RNA-Sequencing | https://www.ncbi.nlm.nih.gov/geo/query/acc.cgi |
| 7. | GSE142699 | *miR-221, miR-222* | Micro-array | https://www.ncbi.nlm.nih.gov/geo/query/acc.cgi?acc=GSE142699 |

**Table 3. Univariable Analysis of *HOTAIRM1*, *miR-222* in AML**

| **Variable** | **Median Survival** | **Hazard Ratio** | **95% Confidence Interval** | **P-value** |
| --- | --- | --- | --- | --- |
| **Event Free Survival** |  |  |  |  |
| ***HOTAIRM1* *Variant 1*** Complete Cohort  AML Favourable  AML Intermediate & Poor  AML t(8;21) | High: 9.85 (n=52)  Low: 9.92 (n=54)  High: 9.99 (n=19)  Low:18.33(n=19)  High: 11.61 (n=22)  Low: 9.16 (n=21)  High: 9.61 (n=16)  Low: 11.70 (n=16) | 0.85  0.98  0.76  1.04 | 0.5508-1.336  0.4477-2.163  0.3896-1.512  0.4562-2.382 | 0.64  0.92  0.95  0.89 |
| ***HOTAIRM1* *Variant 2***  Complete Cohort  AML Favourable    AML Intermediate & Poor    AML t(8;21) | High: 9.52 (n=53)  Low: 9.92 (n=53)   High: 18.33 (n=19)  Low:11.70 (n=19)  High: 12.86 (n=22)  Low: 9.16 (n=22)   High: 18.33 (n=16)  Low: 9.10 (n=16) | 1.09     0.70    0.69     0.82 | 0.7016-1.703  0.3203-1.567    0.3497-1.373  0.3595-1.883 | 0.49  0.26    0.7  0.2 |
| ***miR-222*** Complete Cohort    AML Favourable  AML Intermediate & Poor  AML t(8;21) | High: 8.54(n=53)  Low: 17.24(n=53)  High: 11.08 (n=19)  Low:18.93(n=19)  High: 10.01 (n=22)  Low: 11.80 (n=21)   High: 7.81 (n=16)  Low: 46.13 (n=16) | 1.929  1.12  1.27  3.46 | 1.219-3.053  0.5071-1.289  0.6467-2.508  1.431-8.381 | **0.005**  0.07  0.53  **0.004** |
| ***miR-221*** Complete Cohort  AML Favourable  AML Intermediate & Poor  AML t(8;21) | High:9.23 (n=53)  Low: 11.80 (n=53)  High: 11.08 (n=19)  Low: 18.93 (n=19)  High: 8.94 (n=22)  Low:12.86(n=21)  High: 10.16 (n=16)  Low:18.33 (n=16) | 1.25  1.12  0.69  1.22 | 0.7969-1.968  0.5071-2.476  0.3545-1.363  0.5383-2.771 | 0.27  0.53  0.63  0.48 |
| **Overall Survival** |  |  |  |  |
| ***HOTAIRM1 Variant 1*** Complete Cohort  AML Favourable    AML Intermediate & Poor  AML t(8;21) | High: 12.13 (n=52)  Low: 15.00 (n=54)  High: 46.13 (n=19)  Low: 47.61 (n=19)   High: 15.15 (n=22)  Low: 11.14 (n=22)  High: 19.95 (n=16)  Low: 47.61 (n=16) | 1.07  0.86  0.97  1.07 | 0.6602-1.744  0.3937-2.384  0.4687-2.033  0.4230-2.745 | 0.98  0.51  0.58  0.72 |
| ***HOTAIRM1* *Variant 2*** Complete Cohort  AML Favourable  AML Intermediate & Poor  AML t(8;21) | High: 11.80 (n=39)  Low: 16.22 (n=39)  High: 46.13 (n=19)  Low: 47.61 (n=19)  High: 23.34 (n=22)  Low:11.14 (n=22)  High: 25.26 (n=16)  Low:48.42 (n=16) | 1.35  0.75  0.69  1.04 | 0.8294-2.201  0.3000-1.889  0.3301-1.456  0.4049-2.698 | 0.33  0.37  0.64  0.62 |
| ***miR-222*** Complete Cohort  AML Favourable  AML Intermediate & Poor  AML t(8;21) | High: 9.23 (n=53)  Low: 25.16 (n=53)  High: 11.05 (n=19)  Low: 63.86 (n=19)  High: 11.94(n=22)  Low: 17.24(n=21)  High: 11.05 (n=16)  Low: 63.86 (n=16) | 2.09  2.70  1.441  2.34 | 1.290-3.414  1.104-6.635  0.6984-2.972  0.9070-6.035 | **0.002**  **0.012**  0.48  **0.032** |
| ***miR-221*** Complete Cohort  AML Favourable  AML Intermediate & Poor  AML t(8;21) | High: 18.10 (n=53)  Low:11.01 (n=53)  High: 46.13 (n=19)  Low: 48.42 (n=19)  High: 11.14 (n=22)  Low: 17.24 (n=21)  High: 15.69 (n=16)  Low: 48.60(n=16) | 0.75  1.30  1.43  1.68 | 0.4686-1.224    0.5149-3.286  0.6780-3.03  0.6638-4.276 | 0.5  0.7  0.3  0.2 |

**Table 4: AML Cell Lines and their Molecular and Phenotypic Characteristics**

| **AML Cell Lines** | **Phenotype** | **Translocation** | **Age of Patient** |
| --- | --- | --- | --- |
| THP1 | Monoblast | MLL-AF9 | Paediatric AML (1Y/Male) |
| MOLM-13 | Monoblast | MLL-AF9 | Adult AML (20Y/Male) |
| MOLM-14 | Monoblast | MLL-AF9 | Adult AML (20Y/Male) |
| Kasumi-1 | Myeloblast | RUNX1-RUNX1T1 | Paediatric AML (7Y/Male) |
| KG1 | Myeloblast | FGFR1OP2-FGFR1 | Adult AML (59Y/Male) |
| HL60 | Promyeloblast | - | Adult AML (35Y/Female) |

**Supplementary Figures:**

| **A** |  |
| --- | --- |
| 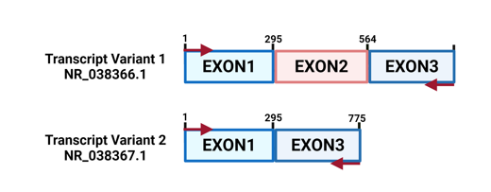  **Primer Design: Cloning HOTAIRM1 Overexpression Construct** | |
| **B** | **C** |
| 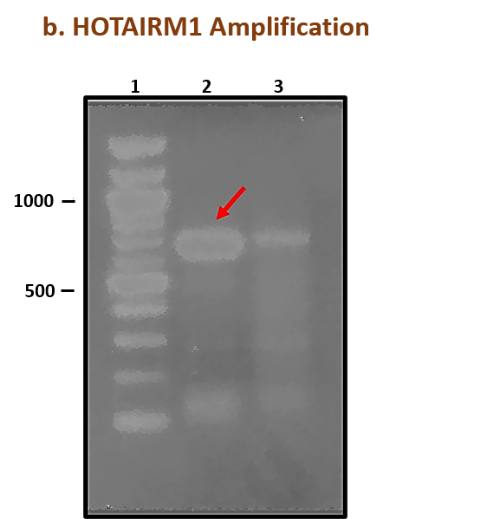 | 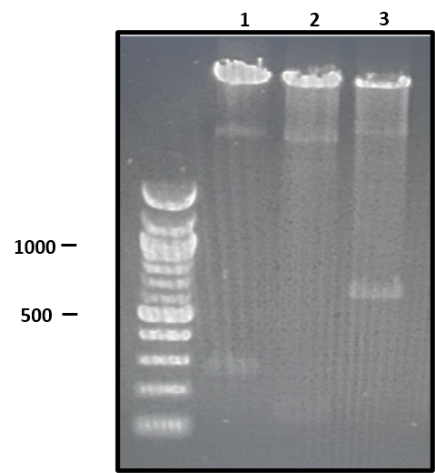 |
| **D** | |
| 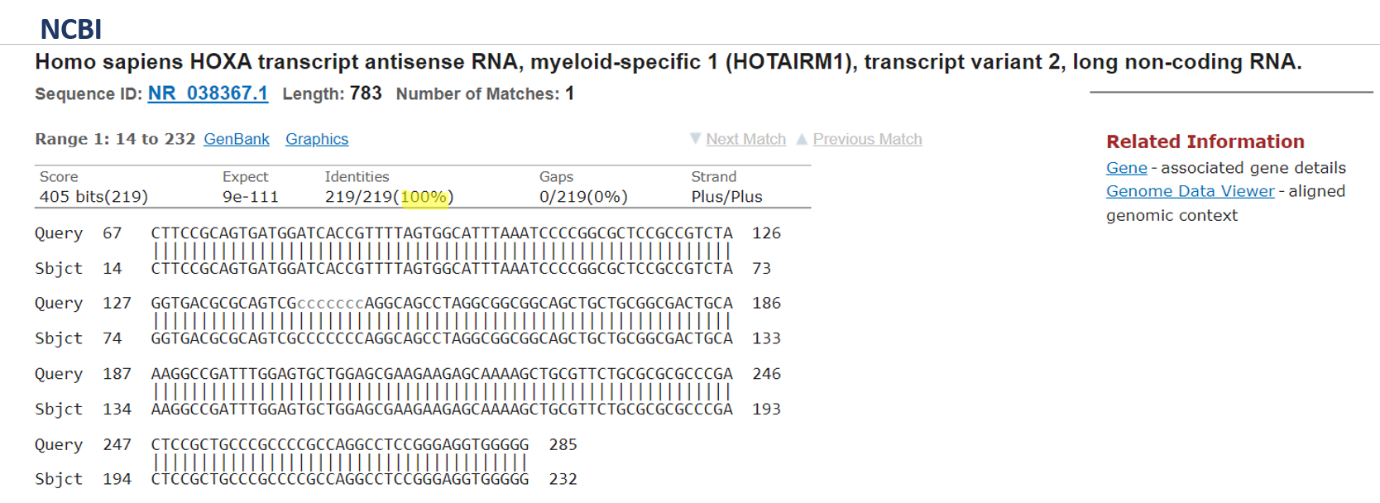 | |
| **E** | |
| 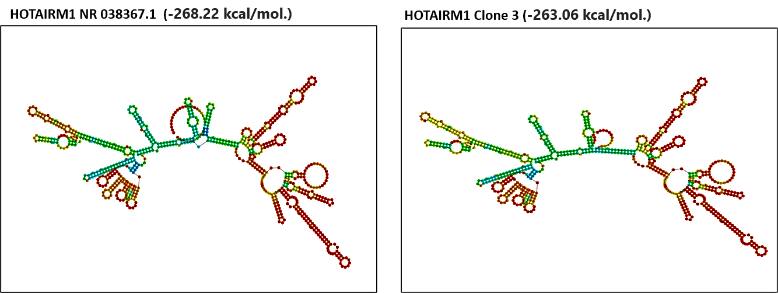 | |
| **F** |  |
| 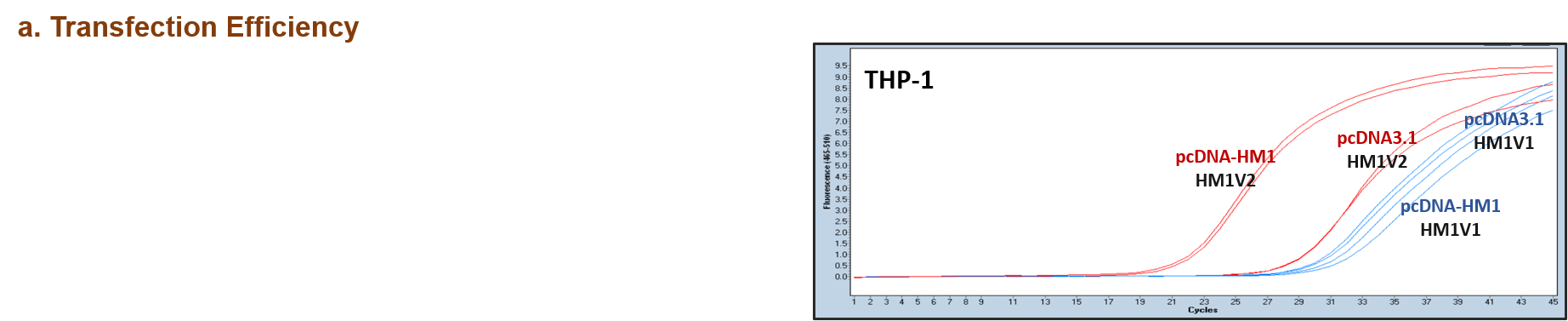 | |
| **G** | |
| 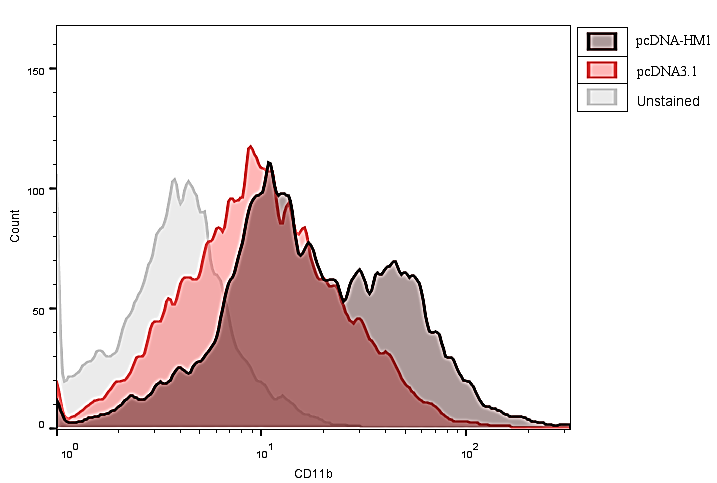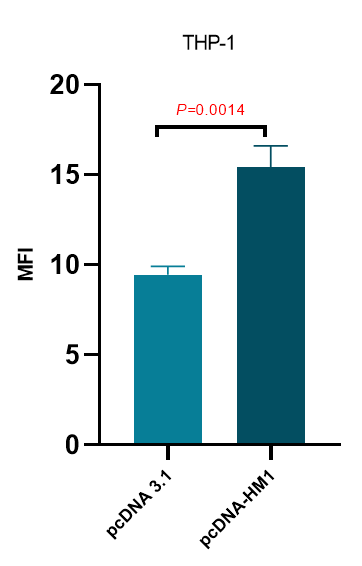 | |
| **H** | |
| 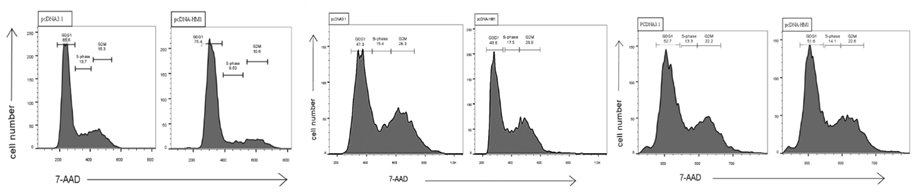  MOLM-13  THP-1  Kasumi-1 | |
| **I** | |
| 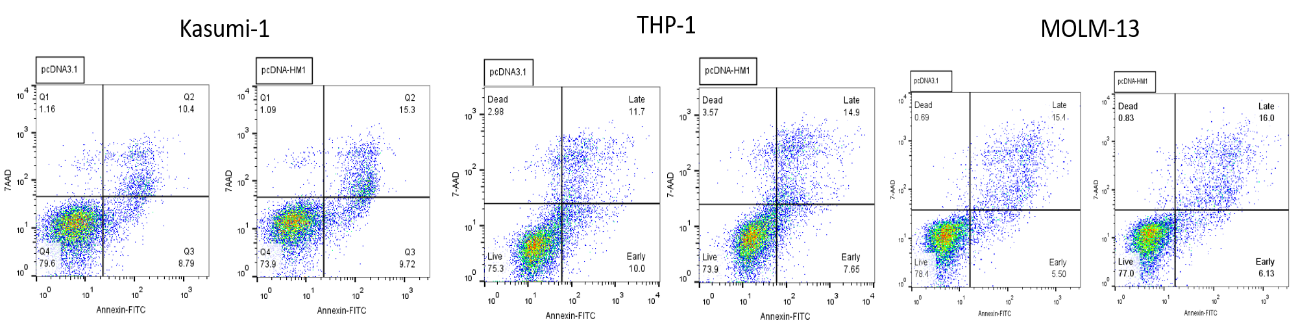 | |
| **J** | |
| 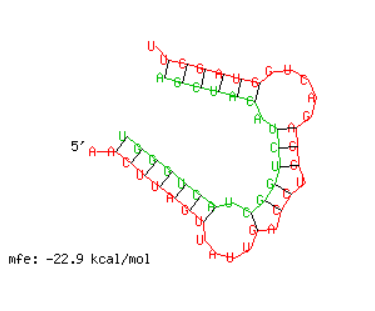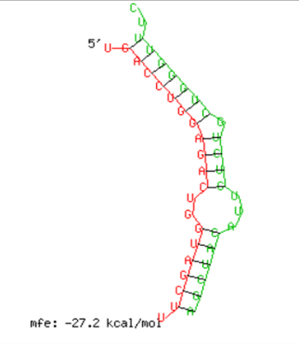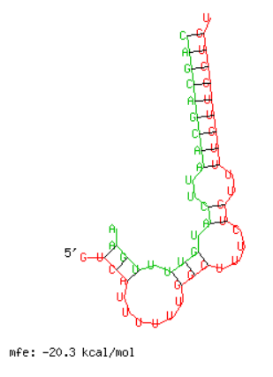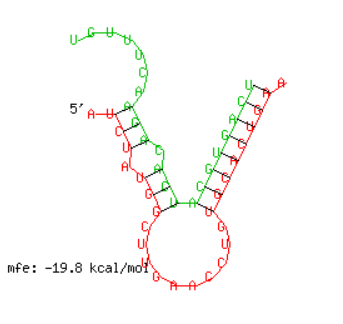  ***HM1V2*-*miR-221***  ***HM1V2*-*miR-222***  ***HM1V2*-*miR-424***  ***HM1V2*-*miR-148b*** | |
| **Supplementary Figure 1A.** Primer design to amplify full length sequence of *HOTAIRM1*. **B.** Gel image of PCR products after amplification of *HOTAIRM1* insert from normal peripheral blood sample. The PCR product amplified was around 700 bp which corresponds to HM1V2 rather than HM1V1. **C.** Gel Image showing results of restriction digestion of clones after ligation. Clone 3 was selected for sequencing as the size is around 700bp which is similar to HM1V2. **D.** NCBI BLAST results show that the sequece obtained after cloning of *HOTAIRM1* is 100% similar to *HOTAIRM1* *variant 2*. **E.** RNAFold generated secondary structure of HOTAIRM1 from NCBI sequence (*HM1V2*) vs. cloned *HM1V2* sequence. **F.** Results from qPCR analysis of *HOTAIRM1* expression in THP-1 cells transfected with pcDNA-HM1 vs. pcDNA3.1 empty vector show that *HM1V2* is overexpressed in transfected cells. **G.** Representative flow cytometry image of increased CD11b expression after transfection of cells with pcDNA-HM1 (dark red), and quantification of MFI shown in the bar graph (right). **H.** Representative image of cell cycle analysis after overexpression of *HOTAIRM1* in Kasumi-1, THP-1 and MOLM-13 cell lines. **I.** Representative image of apoptosis analysis after overexpression of *HOTAIRM1* in Kasumi-1, THP-1 and MOLM-13 cell lines. **J.** Predicted secondary structures of selected microRNA tragets of *HM1V2*. Minimum free energy (mfe) structures were generated using RNAfold. The structures represent HM1V2-miR-148b (mfe: –19.8 kcal/mol), HM1V2-miR-424 (mfe: –20.3 kcal/mol), HM1V2-miR-222 (mfe: –22.9 kcal/mol), and HM1V2-miR-221 (mfe: –27.2 kcal/mol. Red and green colors denote paired and unpaired nucleotides, respectively. | |

| **A** | |
| --- | --- |
| 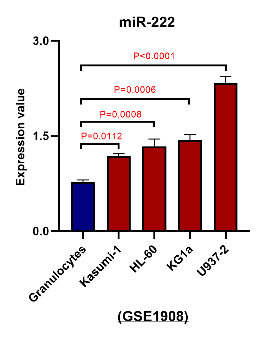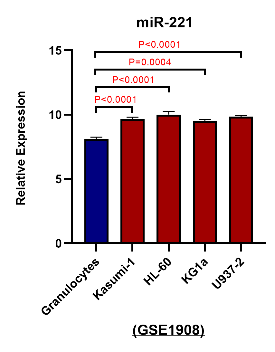  (GSE51908) | |
| **B** | |
| 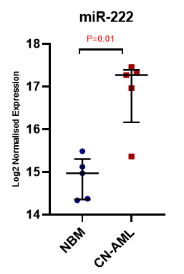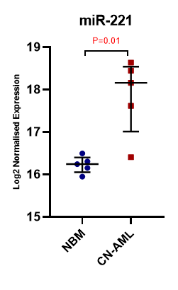 | |
| **C** | **D** |
| 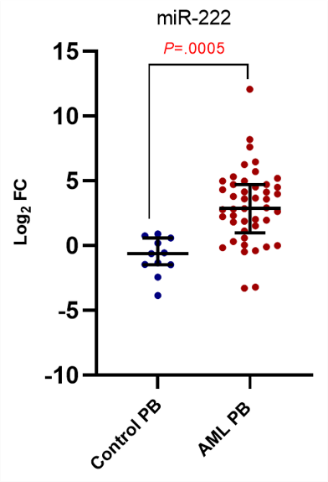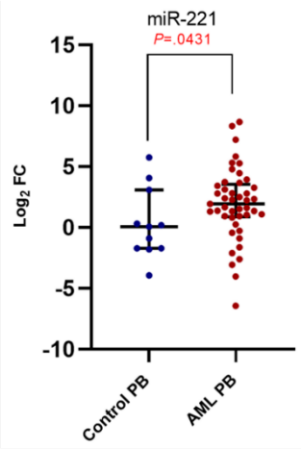 | 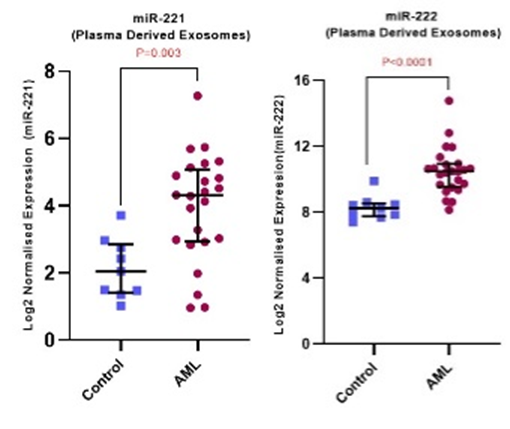  **(GSE142699)** |
| **E** | |
| 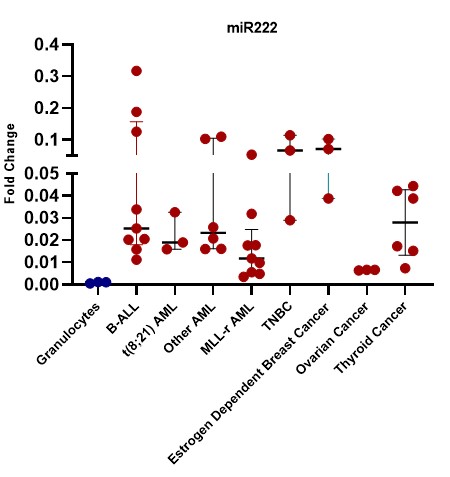 | 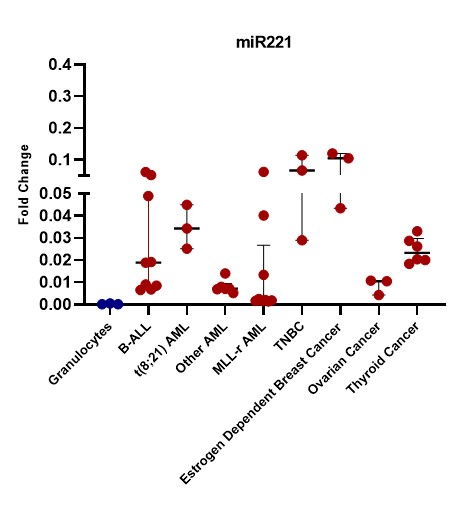 |
| **F** | |
| 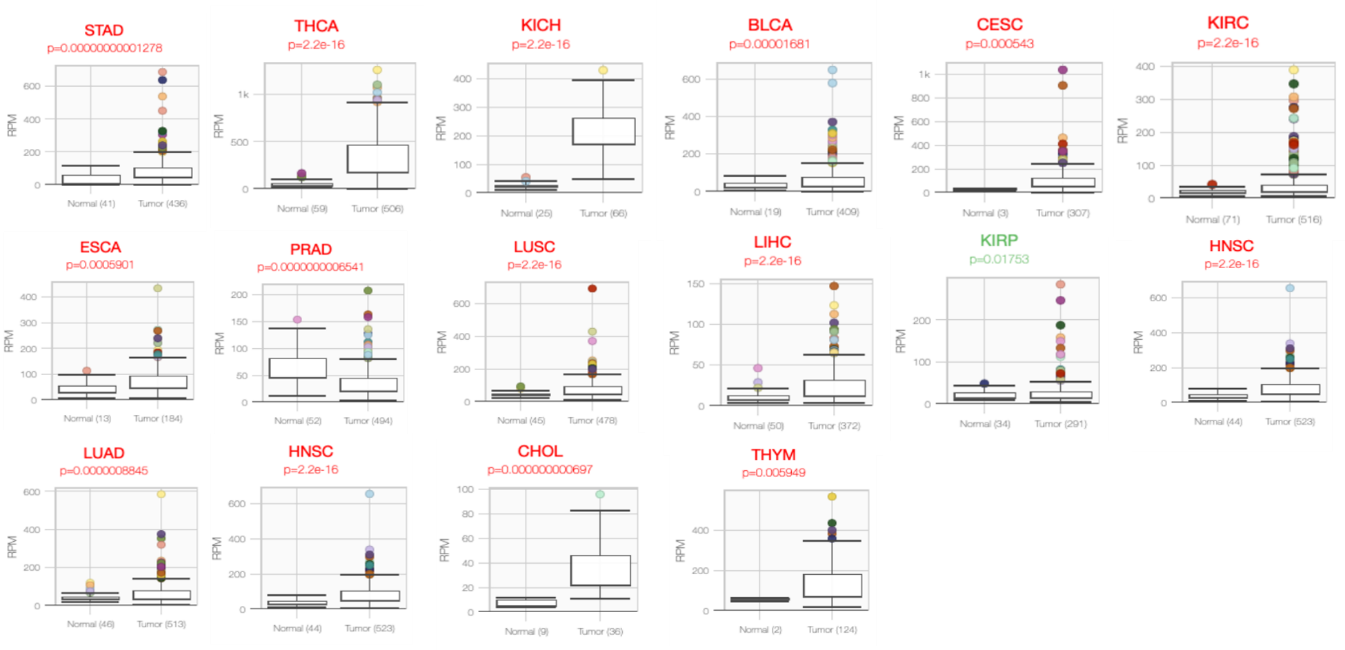  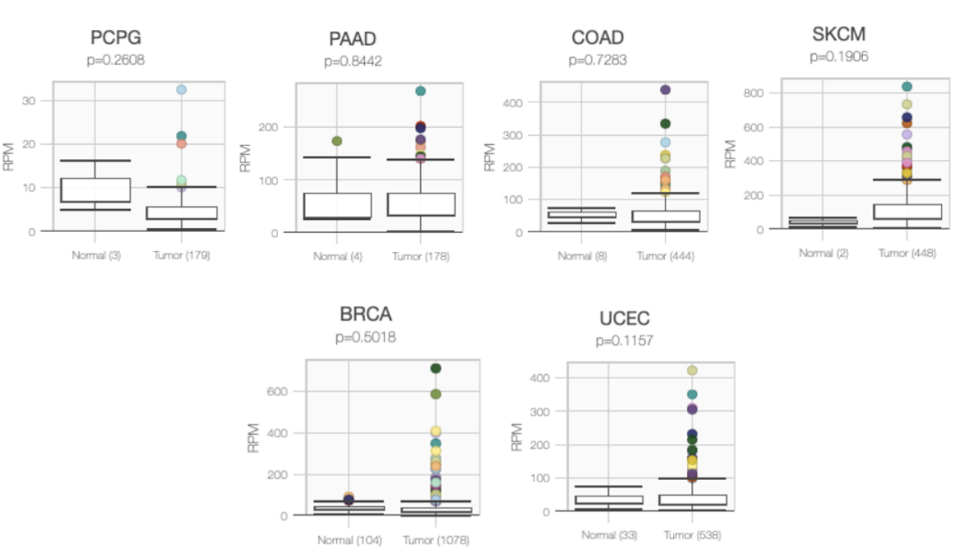 | |
| **G** | |
| 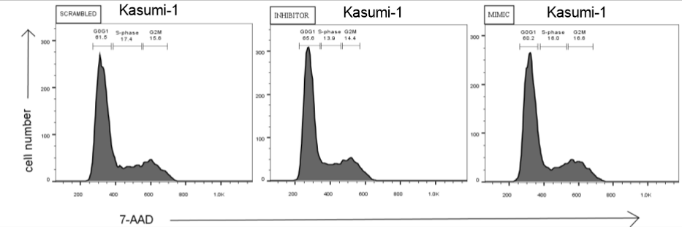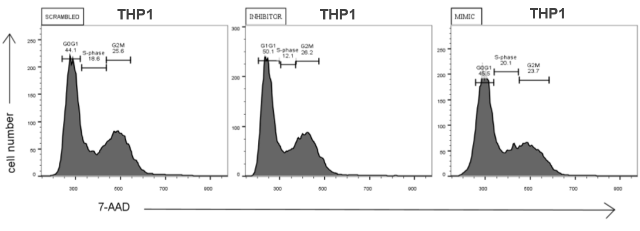 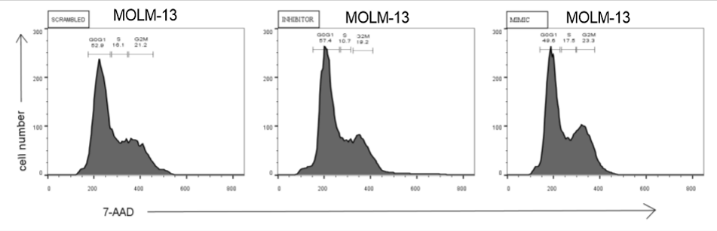 | |
| **H** | **I** |
| 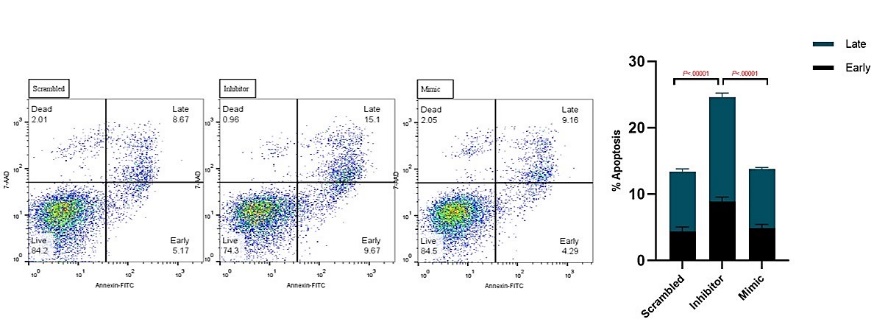**Kasumi-1** | 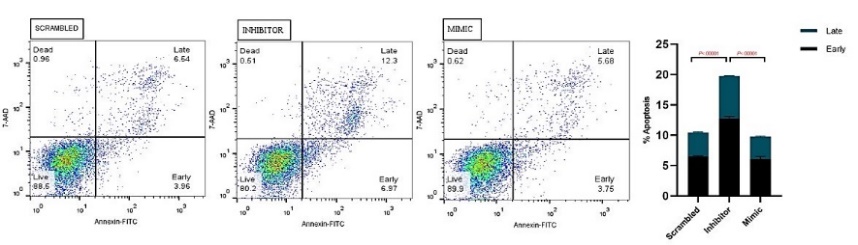**THP-1** |
| **J** |  |
| 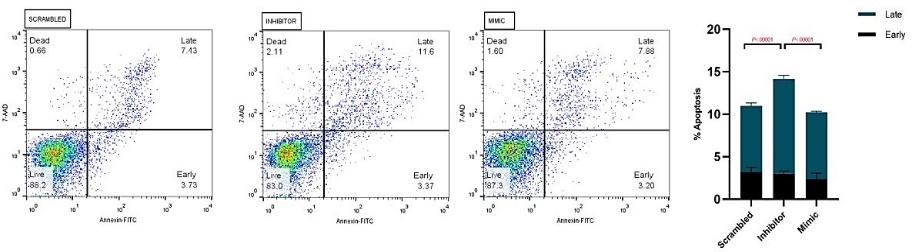**MOLM-13** | |
| **Supplementary Figure 2. Deregulation of miR-222 and 221 in various cancers and regulatory role in AML. A.** Expression of *miR-221* and *miR-222* in AML cell lines, from the GSE51908 dataset. **B.** High expression of *miR-221* and *miR-222* was observed in an independent miR-seq dataset comparing cytogenetically normal (CN-AML) patients with healthy controls. **C.** The high expression of *miR-222* and *miR-221* was validated in PB samples from pediatric patients with AML and found to be significantly higher (P=.0005),(P=0.0431), respectively. **D.** The microRNAs *miR-222* and *miR-221* are detectable in plasma derived exosomes and both microRNAs are significantly upregulated in exosomes from plasma of patients with AML vs controls (GSE142699) (*miR-221* P=0.003, *miR-222* P<0.0001). **E.** Both *miR-222* and *miR-221* are highly expressed in AML, ALL, TNBC, estrogen dependent breast cancer, ovarian cancer and thyroid cancer. **F.** TCGA: The expression of microRNAs was available in 22 TCGA cancer datasets, *miR-222* was significantly upregulated in 16 out of 22 cancer datasets. **G.** Representative images showing that downregulation of *miR-222* in AML cell lines Kasumi-1, THP-1 and MOLM-13 resulted in G0G1 arrest with fewer cells enetering into S-phase. **H,I,J.** Representative images showing that modulation of the expression of *miR-222* in AML cell lines Kasumi-1, THP-1 and MOLM-13 resulted in increased levels of apoptosis. | |

| **A** | **B** | |
| --- | --- | --- |
| 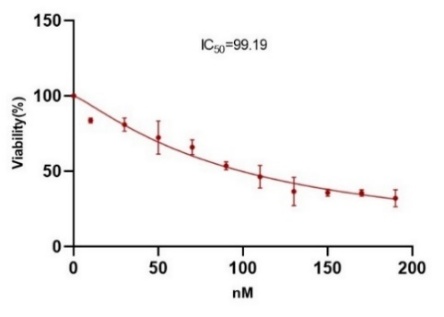 | 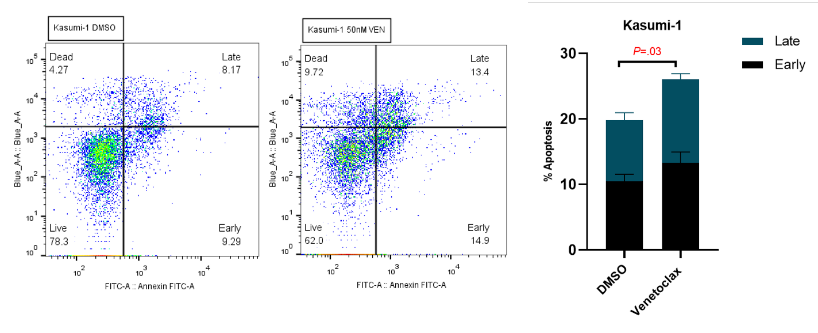 | |
| **C** | | |
| 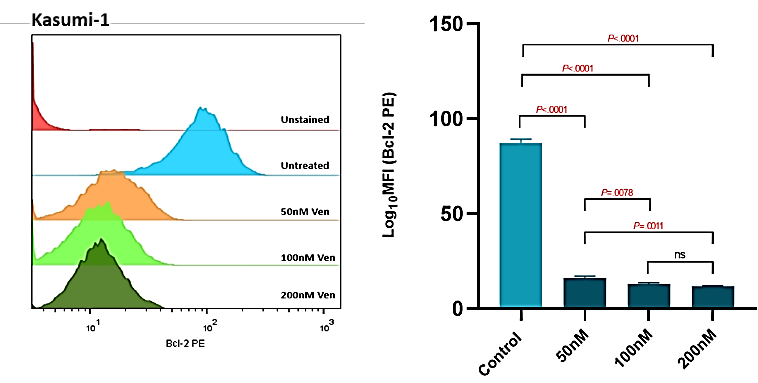 | | |
| **D** | | |
| 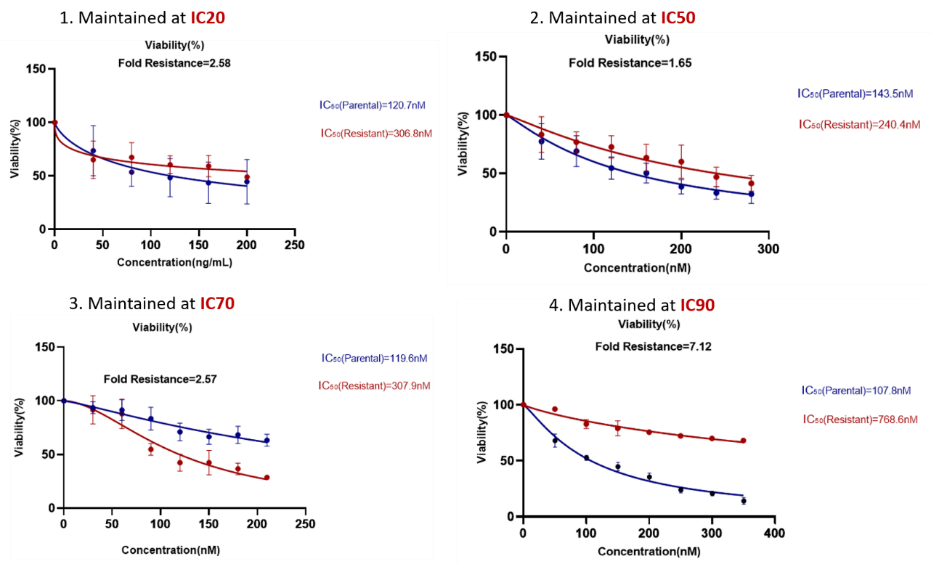 | | |
| **E** | | |
| 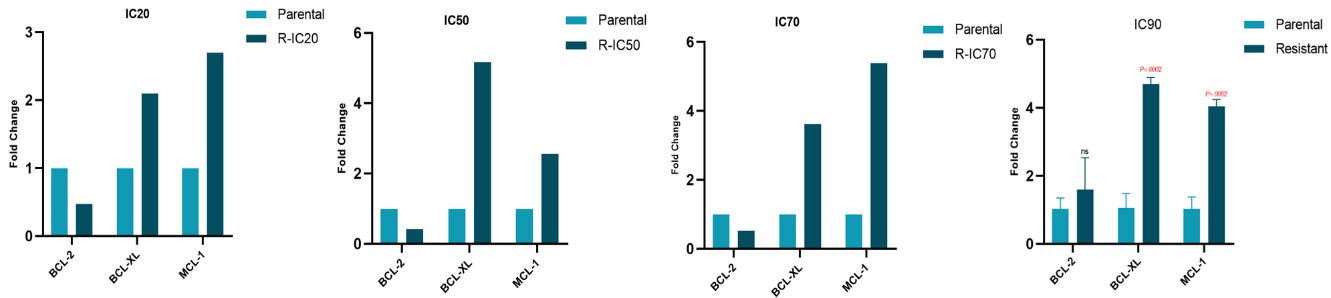 | | |
| **F** | | |
| 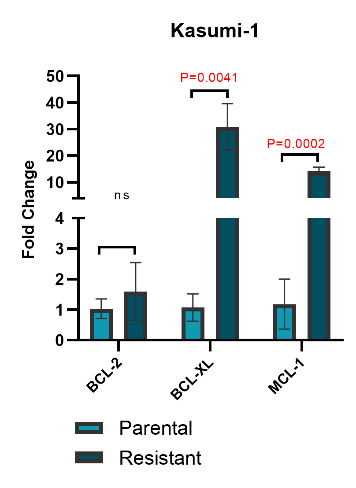 | | |
| **G** | | |
| 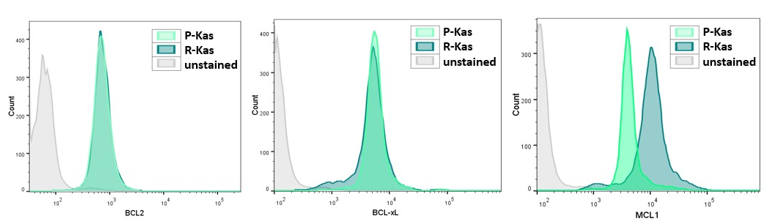 | | |
| 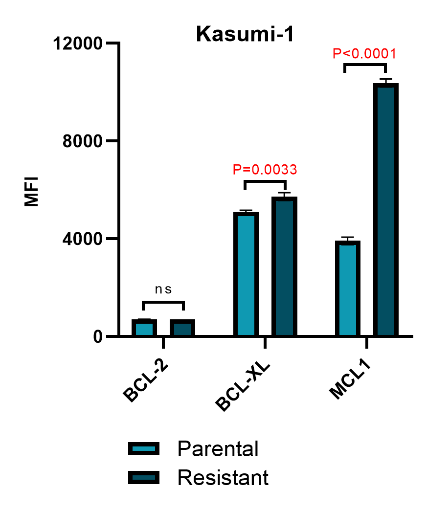 | | |
| **H** | | |
| 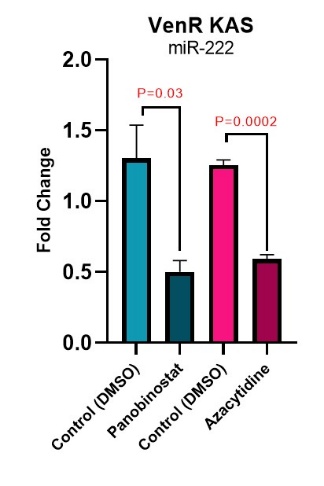 | | |
| **I** | **J** | |
| 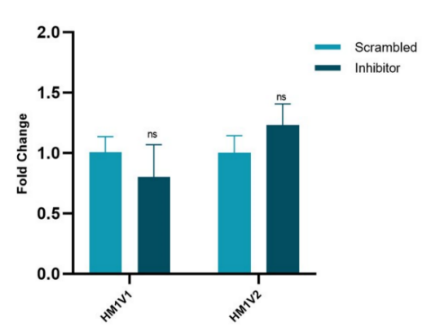 | | 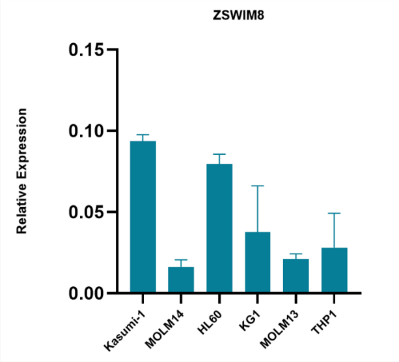 |
| **Supplemetary Figure 3A.** Determination of IC_50_ value of Venetoclax on Kasumi-1 cell line (IC_50_=100nM). **B.** Apoptosis assay using annexin 7-AAD demonstrated that at 50nM concentration of venetoclax (<IC_50_), cells are around 60% viabile. **C.** There is a decrease in protein levels of BCL-2 when venetoclax concentration is increased from 50nM to 100nM (P<0.0001), however, when concentrations are increased to 200nM, the levels of BCL-2 protein doesn’t decrease further. **D.** MTT assay and IC_50_ determination of cells that were maintained in prolonged exposure to venetoclax (at IC20=2.58 fold resistance, IC_50_=1.65 fold resistance, IC70=2.57 fold resistance, IC90=7.12 fold resistance). **E.** Transcript levels of BCL-2, BCL-xL and MCL-1 in cells that were maintained at different concentrations of venetoclax for prolonged periods. Resistant cells at initial IC90 values had increased level of BCL-XL (P=0.0002) and BCL-2 (P=0.0002). **F.** Independent experiement performed in triplicaes. **G.** Protein levels of BCL-2, BCL-XL (P=0.0013) and MCL-1 (P<0.0001) in resistant cells maintained at IC_90_ values vs. sensitive Kasumi-1 cells. **H.** qPCR analysis in VenR KASUMI-1 cells shows significant downregulation of miR-222 following treatment with panobinostat (P = 0.03) and azacytidine (P = 0.0002) compared to DMSO controls**. I.** Downregulation of *miR-222* in Kasumi-1 cell lines does not rescue *HOTAIRM1* expression levels. **J.** ZSWIM8 expression levels are high in t(8;21) AML Kasumi-1 cell line indicating potentially active TDMD pathway. | | |

| **A** | **B** |
| --- | --- |
| 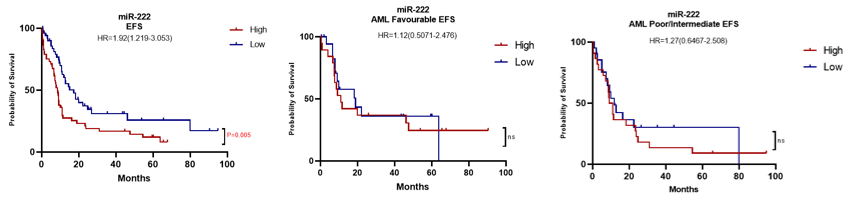 | |
| **C** | **D** |
| 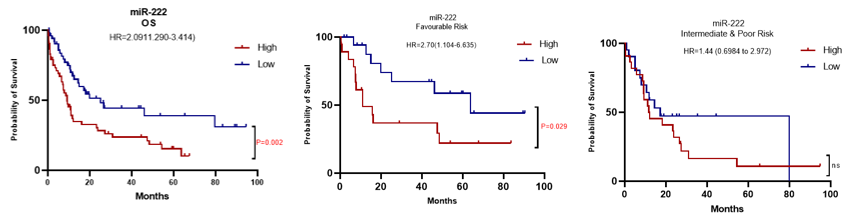 | |
| **E** | |
| 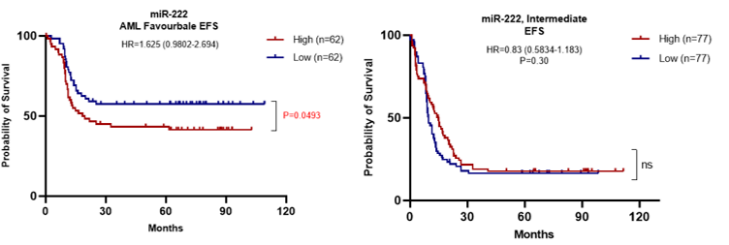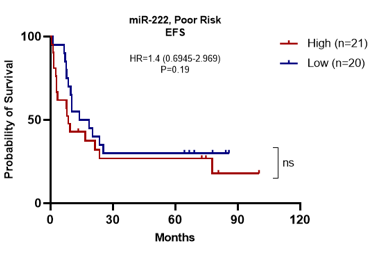 | |
| **F** | |
| 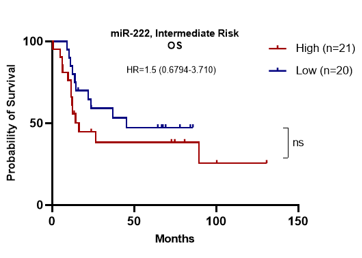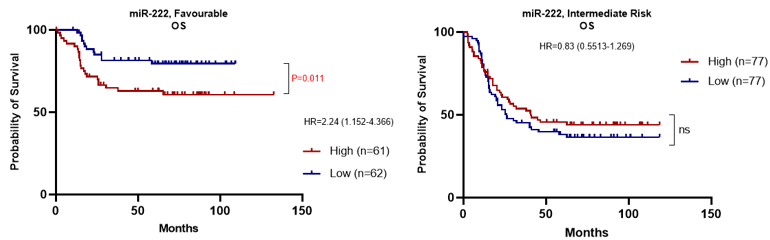 | |
| **Supplemetary Figure 4 A,B.** Event-free survival analysis for patients with high versus low *miR-222* expression in IRCH AML cohorts. **C,D.** Overall survival for patients with high versus low *miR-222* expression in IRCH AML cohorts. **E.** EFS and **O.** OS analysis of *miR-222* high vs low groups in specific patient cytogenetic subgroups i.e. favourable, intermediate and poor (TARGET AML cohort)**.** | |

**Supplementary Code (R 4.4)**

### 1. Sequencing Data Preprocessing

*# Load required libraries*
library(edgeR)
library(DESeq2)
library(limma)
library(tidyverse)

*# Load count matrix and sample metadata*
counts <- read.csv("counts.csv", row.names = 1)
metadata <- read.csv("metadata.csv")

*# Create DESeq2 dataset*
dds <- DESeqDataSetFromMatrix(countData = counts, colData = metadata, design = ~ condition)
 *\# Pre-filtering*
dds <- dds[rowSums(counts(dds)) > 10, ]

*# Normalization*
dds <- estimateSizeFactors(dds)
norm_counts <- counts(dds, normalized=TRUE)

### 2. Differential Expression Analysis

*# Run DESeq2*
dds <- DESeq(dds)

*# Extract results*
res <- results(dds, contrast = c("condition", "AML", "Control"))

*# Shrinkage estimation for fold changes*
resLFC <- lfcShrink(dds, coef=2, type="apeglm")

*# Save results*
write.csv(as.data.frame (resLFC), "DEG_results.csv")

**3. Multivariable Analysis**

**1. Event Free Survival**
*#Load libraries*

library(survival)

library(survminer)

library(readxl)

library(writexl)

*# Load the data*

data <- read_excel("D:/AIIMS/PhD Thesis/Experiments/All Experiments compilation/Data compilation for thesis/Clinical analysis/IRCH/Multivariable/MultivariableEFS - Favourable.xlsx")

*# Display the first few rows and column names to check for any inconsistencies*

print(head(data))

print(colnames(data))

*# Convert necessary columns to numeric if not already*

data$age_yrs <- as.numeric(data$age_yrs)

data$hb <- as.numeric(data$hb)

data$platelet <- as.numeric(data$platelet)

data$tlc <- as.numeric(data$tlc)

data$pblast <- as.numeric(data$pblast)

data$bmblast <- as.numeric(data$bmblast)

data$miR222 <- as.numeric(data$miR222)

data$Months_EFS <- as.numeric(data$Months_EFS)

data$Event<- as.numeric(data$Event)

*# Create binary variables based on median split for continuous variables*

data$age_yrs_above_median <- ifelse(data$age_yrs > median(data$age_yrs, na.rm = TRUE), 1, 0)

data$hb_above_median <- ifelse(data$hb > median(data$hb, na.rm = TRUE), 1, 0)

data$platelet_above_median <- ifelse(data$platelet > median(data$platelet, na.rm = TRUE), 1, 0)

data$tlc_above_median <- ifelse(data$tlc > median(data$tlc, na.rm = TRUE), 1, 0)

data$pblast_above_median <- ifelse(data$pblast > median(data$pblast, na.rm = TRUE), 1, 0)

data$bmblast_above_median <- ifelse(data$bmblast > median(data$bmblast, na.rm = TRUE), 1, 0)

data$miR222_high <- ifelse(data$miR222 > median(data$miR222, na.rm = TRUE), 1, 0)

*# Convert categorical variables to factors*

data$sex <- as.factor(data$sex)

data$Risk <- as.factor(data$Risk)

*# Perform Cox proportional hazards regression including all variables*

cox_model <- coxph(Surv(Months_EFS, Event) ~ sex + age_yrs_above_median + hb_above_median +

platelet_above_median + tlc_above_median + pblast_above_median +

bmblast_above_median + Risk + miR222_high, data = data)

*# Summarize the Cox model*

cox_summary <- summary(cox_model)

*# Extract relevant information*

results <- data.frame(

Variable = rownames(cox_summary$coefficients),

HR = round(cox_summary$coefficients[, "exp(coef)"], 2),

CI_Lower = round(cox_summary$conf.int[, "lower .95"], 2),

CI_Upper = round(cox_summary$conf.int[, "upper .95"], 2),

p_value = round(cox_summary$coefficients[, "Pr(>|z|)"], 4)

)

*# Save the results to an Excel file*

write_xlsx(results, path = "cox_regression_results_EFS2_miR222.xlsx")

**2. Overall Survival**

*# Load libraries*

library(survival)

library(survminer)

library(readxl)

library(writexl)

*# Load the data*

data <- read_excel("D:/AIIMS/PhD Thesis/Experiments/All Experiments compilation/Data compilation for thesis/Clinical analysis/IRCH/Multivariable/MultivariableOS - Favourable.xlsx")

*# Display the first few rows and column names to check for any inconsistencies*

print(head(data))

print(colnames(data))

*# Convert necessary columns to numeric if not already*

data$age_yrs <- as.numeric(data$age_yrs)

data$hb <- as.numeric(data$hb)

data$platelet <- as.numeric(data$platelet)

data$tlc <- as.numeric(data$tlc)

data$pblast <- as.numeric(data$pblast)

data$bmblast <- as.numeric(data$bmblast)

data$miR222 <- as.numeric(data$miR222)

data$Months_OS <- as.numeric(data$Months_OS)

data$OS_Event <- as.numeric(data$OS_Event)

*# Create binary variables based on median split for continuous variables*

data$age_yrs_above_median <- ifelse(data$age_yrs > median(data$age_yrs, na.rm = TRUE), 1, 0)

data$hb_above_median <- ifelse(data$hb > median(data$hb, na.rm = TRUE), 1, 0)

data$platelet_above_median <- ifelse(data$platelet > median(data$platelet, na.rm = TRUE), 1, 0)

data$tlc_above_median <- ifelse(data$tlc > median(data$tlc, na.rm = TRUE), 1, 0)

data$pblast_above_median <- ifelse(data$pblast > median(data$pblast, na.rm = TRUE), 1, 0)

data$bmblast_above_median <- ifelse(data$bmblast > median(data$bmblast, na.rm = TRUE), 1, 0)

data$miR222_high <- ifelse(data$miR222 > median(data$miR222, na.rm = TRUE), 1, 0)

*# Convert categorical variables to factors*

data$sex <- as.factor(data$sex)

data$Risk <- as.factor(data$Risk)

*# Perform Cox proportional hazards regression including all variables*

cox_model <- coxph(Surv(Months_OS, OS_Event) ~ sex + age_yrs_above_median + hb_above_median +

platelet_above_median + tlc_above_median + pblast_above_median +

bmblast_above_median + Risk + miR222_high, data = data)

*# Summarize the Cox model*

cox_summary <- summary(cox_model)

*# Extract relevant information*

results <- data.frame(

Variable = rownames(cox_summary$coefficients),

HR = round(cox_summary$coefficients[, "exp(coef)"], 2),

CI_Lower = round(cox_summary$conf.int[, "lower .95"], 2),

CI_Upper = round(cox_summary$conf.int[, "upper .95"], 2),

p_value = round(cox_summary$coefficients[, "Pr(>|z|)"], 4)

)

*# Save the results to an Excel file*

write_xlsx(results, path = "cox_regression_results_OS_miR222.xlsx")

### 3. Machine Learning for Risk Prediction

**1. Random Forest**

*# Load necessary libraries*

library(randomForest)

install.packages("caret")

library(caret)

library(openxlsx) # For exporting to Excel

*# Load your dataset*

data <- read_excel("D:/AIIMS/PhD Thesis/Experiments/All Experiments compilation/Data compilation for thesis/Clinical analysis/TARGET AML/Multivariable/EFS/EFS - t(8;21).xlsx")

*# Convert categorical variables to factors (if applicable)*

data$Gender <- as.factor(data$Gender)

data$FLT3 <- as.factor(data$FLT3)

data$NPM <- as.factor(data$NPM)

data$CEBPA <- as.factor(data$CEBPA)

*# Remove the columns 'TARGET_USI' and 'First_Event' for model training, but keep them in the data*

data_clean <- data[, !(names(data) %in% c("TARGET_USI", "First_Event"))]

*# Step 1: Check for missing values in the dataset*

missing_values <- colSums(is.na(data_clean))

print("Missing values per column:")

print(missing_values)

*# Step 2: Remove rows with missing values (if you want to remove them)*

data_clean <- na.omit(data_clean)

*# Step 3: If you want to impute missing values instead, use this code:*

\# Uncomment the lines below to perform median imputation if preferred

\# preprocessed_data <- preProcess(data_clean, method = 'medianImpute')

\# data_clean <- predict(preprocessed_data, data_clean)

*# Check if any missing values are left*

missing_values_after <- colSums(is.na(data_clean))

print("Missing values after cleaning:")

print(missing_values_after)

*# Set the target variable (EFS_Months) and predictors (excluding the removed columns)*

target_variable <- "EFS_Months"

predictors <- setdiff(names(data_clean), target_variable)

*# Step 4: Create the Random Forest model with the cleaned data*

rf_model <- randomForest(EFS_Months ~ ., data = data_clean[, c(predictors, target_variable)])

*# Print the model summary*

print(rf_model)

*# Step 5: Get the feature importance scores*

importance_scores <- rf_model$importance

*# View feature importance*

print("Feature Importance Scores:")

print(importance_scores)

*# Step 6: Plot the feature importance*

varImpPlot(rf_model)

*# Step 7: Export the full dataset (with 'TARGET_USI' and 'First_Event') to an Excel file*

write.xlsx(data, "path/to/your/output.xlsx") # replace with your desired output file path

**2. XGboost**

*# Load necessary libraries*

library(readxl)

library(openxlsx)

library(xgboost)

library(caret)

library(dplyr)

*# Load dataset*

data <- read_excel("D:/AIIMS/PhD Thesis/Experiments/All Experiments compilation/Data compilation for thesis/Clinical analysis/TARGET AML/Multivariable/EFS/EFS - t(8;21).xlsx")

*# Convert categorical variables to factors*

data$Gender <- as.factor(data$Gender)

data$FLT3 <- as.factor(data$FLT3)

data$NPM <- as.factor(data$NPM)

data$CEBPA <- as.factor(data$CEBPA)

*# Remove non-predictor columns for model training*

data_clean <- data[, !(names(data) %in% c("TARGET_USI", "First_Event"))]

\# Remove rows with missing values (or use imputation)

data_clean <- na.omit(data_clean)

\# Convert factor variables to numeric (required for xgboost)

data_clean[] <- lapply(data_clean, function(x) {

if (is.factor(x)) as.numeric(as.factor(x)) else x

})

*# Set target and predictor variables*

target_variable <- "EFS_Months"

predictors <- setdiff(names(data_clean), target_variable)

*# Prepare matrices for xgboost*

xgb_matrix <- xgb.DMatrix(data = as.matrix(data_clean[, predictors]),

label = data_clean[[target_variable]])

*# Set parameters for regression*

params <- list(

objective = "reg:squarederror",

eval_metric = "rmse"

)

*# Train the model*

xgb_model <- xgb.train(

params = params,

data = xgb_matrix,

nrounds = 100,

verbose = 1

)

*# View model info*

print(xgb_model)

*# Feature importance*

importance <- xgb.importance(model = xgb_model, feature_names = predictors)

print(importance)

*# Plot feature importance*

xgb.plot.importance(importance_matrix = importance)
